# Distinct differentiation trajectories leave lasting impacts on gene regulation and function of V2a interneurons

**DOI:** 10.1101/2024.12.03.626573

**Authors:** Nicholas H. Elder, Alireza Majd, Emily A. Bulger, Ryan M. Samuel, Lyandysha V. Zholudeva, Todd C. McDevitt, Faranak Fattahi

## Abstract

During development, early regionalization segregates lineages and directs diverse cell fates. Sometimes, however, distinct progenitors produce analogous cell types. For example, V2a neurons, are excitatory interneurons that emerge from different anteroposterior progenitors. V2a neurons demonstrate remarkable plasticity after spinal cord injury and improve motor function, showing potential for cell therapy. To examine how lineage origins shape their properties, we differentiated V2a neurons from hPSC-derived progenitors with distinct anteroposterior identities. Single-nucleus multiomic analysis revealed lineage-specific transcription factor motifs and numerous differentially expressed genes related to axon growth and calcium handling. Bypassing lineage patterning via transcription factor-induced differentiation yielded neurons distinct from both developmentally relevant populations and human tissue, emphasizing the need to follow developmental steps to generate authentic cell identities. Using *in silico* and *in vitro* loss-of-function analyses, we identified CREB5 and TCF7L2 as regulators specific to posterior identities, underscoring the critical role of lieage origins in determining cell states and functions.

## Introduction

Human pluripotent stem cell (hPSC)-derived tissues offer immense potential for disease modeling and novel therapeutic interventions. A major question in the field has been how to best transition pluripotent stem cells into desired cell lineages. This endeavor has led to methods that either mimic developmental processes through ‘directed differentiations’ or rely on transcription factor overexpression in ‘induced differentiations’. During development, temporally distinct and spatially separated progenitor populations may give rise to seemingly analogous cell types with common morphological features and expression of hallmark genes. However, these lineally separated populations often differ tissue localization or functional capacity. Some examples of this include cardiomyocytes arising from the first or second heart fields, melanocytes differentiated from neural crest or Schwann cell-derived precursors (Samuel et al., 2023), and neurons of the central and peripheral nervous systems (Alshawaf et al., 2018; Fattahi et al., 2016).

The molecular and functional differences that exist between such analogous cell types can be even larger when comparing cells generated using developmentally relevant strategies to those generated through induced differentiation via transcription factor over expression in stem cells (Elder et al., 2022; Fernandopulle et al., 2018; Zhang et al., 2013). Induced cells can express key marker genes and morphologically resemble the target population, but by nature of having skipped developmental stages, can also misregulate transcription and chromatin marks. These differences necessitate the use of -omics level approaches to deeply characterize the transcriptional and chromatin landscape of cell populations and to facilitate in-depth comparisons between primary cell populations and those generated *in vitro*.

In this study, we investigate how differentiation trajectories influence the properties of V2a neurons—ventral, excitatory, ipsilaterally-projecting interneurons that emerge along the anteroposterior axis and play key roles in phrenic and locomotor circuits (Ampatzis et al., 2014; Azim et al., 2014; Bubnys et al., 2019; Crone et al., 2012; Hayashi et al., 2018). V2a neurons exhibit plasticity in restoring diaphragm function and are a key target population for the recovery of locomotion via electrical stimulation after spinal cord injury (Kathe et al., 2022; Squair et al., 2023; Zholudeva et al., 2017). Transplantation of V2a neurons also has potential as a cell therapy to enhance functional recovery after spinal cord injury, making this cell type of high therapeutic interest (Bonner & Steward, 2015; Zholudeva et al., 2018; Zholudeva & Lane, 2019).

V2a neurons develop over the entire axial range of the neural tube, and understanding how axial positioning intersects with their identity as a V2a neuron is critical for deriving therapeutically relevant cell types *in vitro* (Hayashi et al., 2018; Menelaou et al., 2014; Menelaou & McLean, 2019). *In vivo* and *in vitro*, axial patterning is one of the first steps of development, preceding much of differentiation (Metzis et al., 2018). Anterior V2a neurons primarily arise from neurectoderm progenitors (NEPs), which are induced by the inhibition of BMP and TGF signaling in pluripotent stem cells. By contrast, posterior V2a neurons are formed from a distinct population termed neuromesodermal progenitors (NMPs) which contribute to the posterior elongation of the embryo as they differentiate into the neural tube and adjacent somites (Cambray & Wilson, 2007; Henrique et al., 2015; Tzouanacou et al., 2009; Wymeersch et al., 2019). Despite differences in early development, subsequent dorsoventral patterning of the neural tube is remarkably conserved along the anteroposterior axis, producing V2a neurons that express the canonical marker VSX2 (also known as CHX10) (Clovis et al., 2016; Debrulle et al., 2019). However, the anteroposterior origin of V2a neurons could lead to molecular differences with potential functional implications. Indeed, V2a neurons exhibit different patterns of axon extension as well as long-term expression of the V2a marker VSX2 as a function of their location within the spinal cord in model organisms (Hayashi et al., 2018; Menelaou et al., 2014).

Here, we interrogate how early differences in developmental patterning and progenitor identity impact the resulting V2a phenotype using three distinct models of hPSC differentiation. We first established a novel system to generate V2a neurons from NMPs to compare their molecular and functional features with their NEP-derived counterparts. We then used multiomic single nucleus RNA and ATAC-seq approaches to examine how differences in the early developmental lineage are imprinted into the chromatin landscape and transcriptional signature of these neurons. We found that while both NEP-derived and NMP-derived V2a populations were similar to human tissue derived V2a neurons, the two *in vitro*-derived populations exhibited notable differences in gene expression and regions of open chromatin stemming from their distinct progenitor origins. Functional profiling of the neurons supported the differences observed in multiomics data, showing NMP-derived populations established network-wide activity more rapidly and consistently than NEP-derived neurons.

To further define the impact of differentiation trajectories on V2a neurons, we also established a protocol for induced differentiation which skips normal developmental patterning and rapidly transitions stem cells to post-mitotic neurons. We find that induced VSX2-expressing neurons greatly differ from both of our NEP and NMP-derived V2a populations, express genes not typically associated with V2a neurons, and shift their identity even after progressing to a post-mitotic state. Lastly, leveraging our multiomics dataset, we employed *in silico* gene regulatory network (GRN) modeling and perturbation to show how different transcription factors regulate lineage-specific differences between NEP and NMP-derived V2a populations. Our results highlight the significance of developmentally accurate directed differentiation strategies to generate relevant and authentic cell types for developmental and disease modeling and therapeutic application.

## Results

Distinct neural progenitor populations can be differentiated to V2a neurons To better understand how the axial identity influences mature V2a cell identity, we began by deriving NEPs and NMPs using previously established protocols before maturing these progenitor populations to V2a neurons under identical conditions (Butts et al., 2017, 2019; Lippmann et al., 2015). Mirroring the conditions of *in vivo* development, hPSCs are converted into NEPs via dual SMAD inhibition in FGF-containing stem cell media for the first 5 days (**Figure 1A**; top method). NMPs are induced *in vivo* and *in vitro* by balanced FGF and WNT signaling and are identified by co-expression of SOX2 and TBXT (also known as Brachyury), as well as the caudal homeodomain protein CDX2. We performed an NMP induction protocol (**Figure 1A**; bottom method) (Lippmann et al., 2015) that resulted in high levels of SOX2 and TBXT coexpression and >90% CDX2 expression at day 5 (**Figure 1B-D**). Notably, TBXT and CDX2 were completely absent in the NEP condition at a matched time point (**Figure 1B-D**).

**Figure 1.**
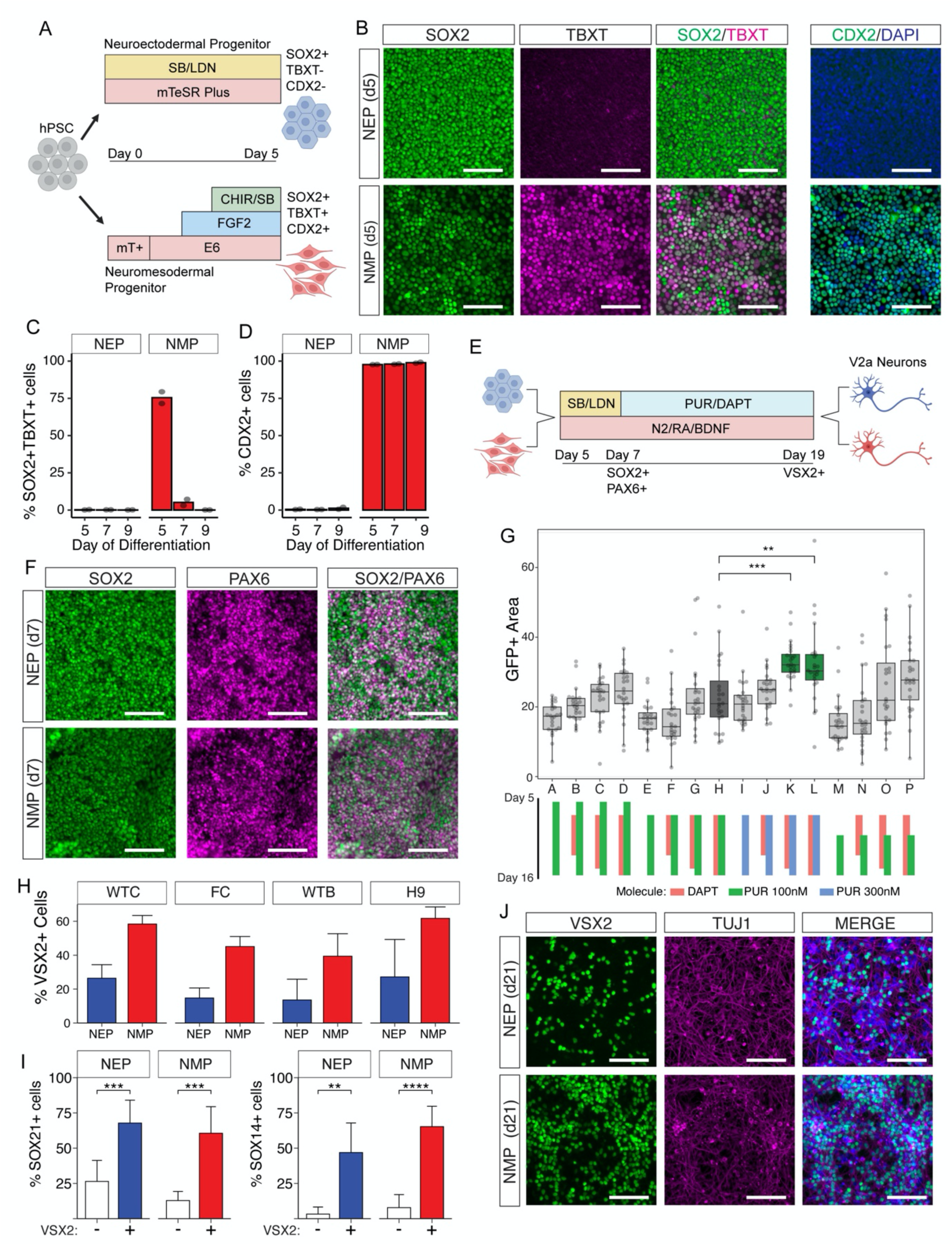
Generation of V2a neurons from distinct progenitor lineages. A. Schematic of the first five days of differentiation from human pluripotent stem cells toward neural progenitors under defined media conditions with expected marker gene expression. Abbreviations: SB = SB431542, LDN = LDN193189, CHIR = CHIR99021, E6= Essential 6, mT+ = mTeSR Plus. B. Representative expression of neuromesodermal progenitor population markers SOX2, TBXT, and CDX2 at day 5 of differentiation in the neuroectodermal and neuromesodermal routes (top and bottom rows, respectively). Note the overlap of SOX2 and TBXT expression in the NMP condition. Scale bars represent 100 um. C. Flow cytometry quantification of the percent of cells coexpressing NMP markers SOX2 and TBXT in both progenitor populations at days 5, 7, and 9 of differentiation. n = 2 biological replicates. D. Flow cytometry quantification of the percent of cells expressing NMP marker CDX2 in both progenitor populations at days 5, 7 and 9 of differentiation. n = 2 biological replicates. E. Schematic of the differentiation of distinct progenitor lineages toward V2a neurons under identical defined media conditions from day 5 to day 19 of differentiation with expected marker gene expression. Abbreviations: SB = SB431542, LDN = LDN193189, PUR = purmorphamine, RA = retinoic acid. F. Representative expression of SOX2 and PAX6 in the NEP and NMP conditions (top and bottom rows, respectively) at day 7 of differentiation. Scale bars represent 100 um. G. Differentiation optimization using the FOXN4:Cre reporter line. Boxplots show the percent of the GFP+ area, as reported by FOXN4:Cre expression, under modified addition of PUR and DAPT between days 5 and 16 of differentiation as indicated by the colored bars (bottom). H (dark gray) represents prior published conditions. K,L (green) show significantly increased GFP+ area with 300 nM PUR. Condition L was selected for further experiments. Abbreviations: PUR = purmorphamine. H. Flow cytometry quantification of the percent of VSX2+ cells in dissociated cultures at day 19 of differentiation from either NEP or NMP lineages in four human stem cell lines. Abbreviations: WTC = WTC11, FC = FOXN4:Cre. n >=2 biological replicates. Error bars represent standard deviation. I. Flow cytometry quantification of the percent of SOX21+ or SOX14+ cells in both VSX2 positive and negative populations at day 19 of differentiation. n = 7 biological replicates. ** is p<0.01, *** is p<0.001, **** is p<0.0001, error bars represent standard deviation. J. Representative expression of VSX2 and TUJ1 in NEP and NMP-derived neurons at day 21, two days after replating. Scale bars represent 100 um.

Following NEP or NMP induction over the first 5 days of differentiation, the progenitor populations were subsequently differentiated toward V2a interneurons under identical media formulations (**Figure 1E**). Since the neural tube patterning is highly conserved between anterior and posterior regions *in vivo*, identical treatment from day 5 onward allowed us to specifically investigate the impact of early progenitor origins on the resulting neurons. We compared the differentiated populations at day 7 following 48 hours of dual SMAD inhibition in the presence of 100 nM RA, a potent neuralizing morphogen. At this time-point, both populations began to co-express canonical neural progenitor markers SOX2 and PAX6, while TBXT levels in the NMP-derived population dropped significantly (**Figure 1C,F**). CDX2 remained highly expressed in the NMP-derived progenitors (**Figure 1D**). These data show the efficient generation of neural progenitors via two different differentiation routes that converge on a SOX2+ PAX6+ neural phenotype.

Prior studies have generated V2a neurons from neural progenitors via balanced control over three developmental signaling pathways: retinoic acid, Sonic hedgehog (SHH), and Notch (Butts et al., 2017, 2019). Retinoic acid generated by the paraxial somites and SHH secreted by the floor plate serve to pattern the neural tube, while the Notch pathway controls the balanced acquisition of excitatory V2a and inhibitory V2b fates from p2 progenitors. *In vitro*, the small molecule agonist purmorphamine (PUR) is substituted for SHH while the gamma-secretase inhibitor DAPT potently inhibits Notch signaling and biases p2 progenitors toward the V2a fate at the expense of the V2b fate. To further optimize the timing and concentration of PUR and DAPT to yield increased V2a neurons in our differentiations, we generated a Cre lineage reporter under the control of the transcription factor FOXN4, which is expressed solely in p2 progenitors. This lineage reporter (called FOXN4:Cre or abbreviated FC) allowed us to observe the efficiency of p2/V2a differentiation in live cultures through the readout of a GFP reporter. We used the FOXN4:Cre line to test 16 iterations of the differentiation protocol where we varied the timing or concentration of PUR and DAPT (**Figure 1G**). We found that 300 nM PUR added from day 7 onward significantly increased the GFP+ area over our control condition with 100 nM PUR (**Figure 1G**). Therefore, this modification was used throughout the rest of this study.

On day 19 of V2a differentiation, cells were dissociated and assessed for the canonical V2a marker VSX2 via flow cytometry (**Figure 1H**). While VSX2+ V2a neurons were detected in all differentiations across four independent cell lines, the NMP condition consistently produced higher percentages of V2a neurons under our experimental conditions (**Figure 1H**). We further validated V2a identity by flow cytometry for secondary markers SOX14 and SOX21 and observed significant enrichment of each marker in the VSX2+ population (**Figure 1I**). Finally, cells that were replated and allowed to grow for an additional two days generated extensive networks of neurites, supporting neural identity (**Figure 1J**). These results confirm the V2a neuron identity in both the NEP- and NMP-derived cultures.

### Differentiated neurons from distinct progenitor lineages recapitulate the identities of the ventral neural tube

Having established neural populations with a canonical V2a identity (VSX2+/SOX14+/SOX21+) via both differentiation methods and across multiple cell lines, we examined whether epigenetic and gene regulatory patterns differed based on early progenitor lineage. To address these questions, we performed single nucleus multiome sequencing after 19 days of differentiation (**Figure 2A**).

**Figure 2.**
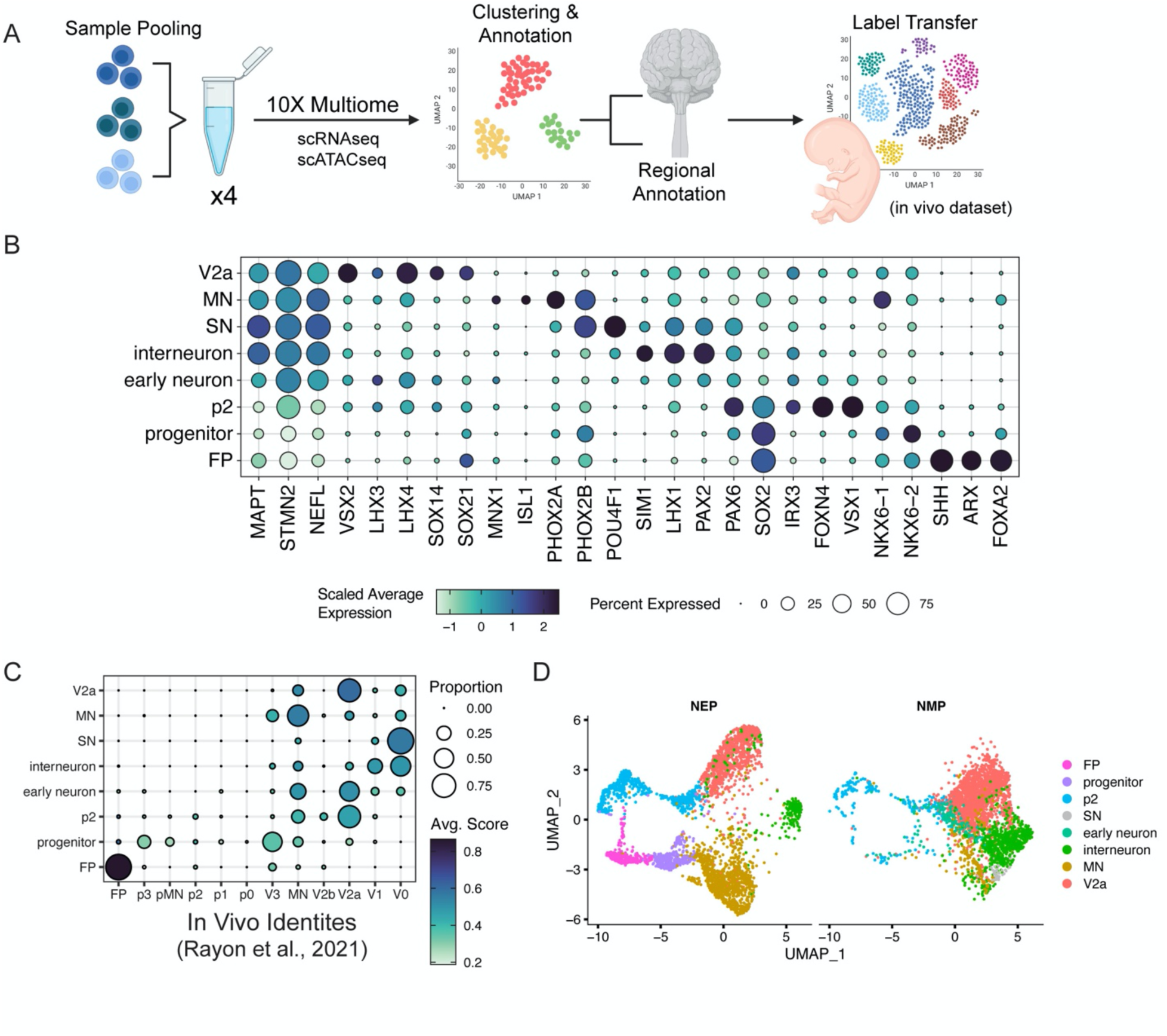
Distinct progenitor lineages recapitulate the cellular diversity of primary human neural tube. A. Schematic of sample pooling for multiome and cell annotation steps. B. Expression of selected marker genes for the 8 annotated cell types in a merged object of all of multiome samples. C. Results of an unbiased label transfer of multiome samples onto ventral human fetal spinal cord cells. D. UMAP of annotated multiome samples after Harmony integration.

We first used the snRNA-seq data to cluster and annotate cell types according to known markers (**Figure 2B, Figure S1)**. V2a neurons expressed VSX2, LHX3, LHX4, SOX21, and SOX14. Motor neurons (MN) were identified by PHOX2A, PHOX2B, and ISL1 expression or MNX1 (also known as HB9) and ISL1 expression, representing cranial or spinal identities, respectively. Overlap of PHOX2B and POU4F1, detected in one NMP-derived sample, was suggestive of a sensory neuron population (SN), possibly arising from a small population of neural crest cells arising during NMP differentiation (**Figure S1G,H, Figure S2A**). Non-V2a ventral interneurons, such as V3, V1, and V0 neurons clustered together as seen by expression of a mix of ventral markers, including SIM1, LHX1, and PAX2 were labeled ‘interneurons’. We also identified a large population of p2 progenitors that expressed PAX6, IRX3, FOXN4, and VSX1. Other cells expressed primarily SOX2 and NKX6.2 which are widely expressed across the ventral progenitor domains and were therefore broadly termed ‘progenitors’. Similarly, a population that expressed neural genes STMN2, NEFL, and MAPT but lacked specific population markers were called ‘early neurons’. Lastly, highly specific expression of SHH, FOXA2, and ARX indicated the presence of floor plate (FP) cells.

When comparing the relative proportions of each cell type in each sample, we again noted that the NMP condition produced a higher proportion of V2a neurons (**Figure S2A**). Only the NEP condition, however, contained floor plate populations. There was also a noticeable shift in the proportion of motor neurons and interneurons between the NEP and NMP conditions. Although both conditions were treated with the same concentration and duration of PUR to produce ventral identities, there is a stronger ventral bias in the identities in the NEP-derived samples compared to those of the NMP condition. This ventral shift is seen in the presence of the floor plate and proportionally larger motor neuron populations, which are ventral to the p2/V2 domain *in vivo* (**Figure S2A**).

We also examined which anteroposterior markers were expressed in our NEP and NMP conditions (**Figure S2B**). We observed low expression of off-target forebrain, midbrain, and eye markers such as FOXG1, OTX2, and CRX (**Figure S2B**). In the NEP-condition, HOX expression primarily ranged from HOX 1-5 paralogs, analogous to a hindbrain identity. NMP-derived samples also expressed these relatively anterior paralogs in addition to more caudal HOX genes such as HOXC5, HOXC6, and HOXB9, consistent with a cervical spinal cord identity (**Figure S2B**). This assessment of axial markers supports a model where the NEP-derived V2a neurons more closely resemble hindbrain populations while NMP-derived neurons resemble spinal populations

To further confirm our assigned cell identities, we performed an unbiased single cell label transfer between our cells and the ventral spinal cell types annotated in an *in vivo* human fetal dataset (Rayon et al., 2021). Label transfer results largely agreed with our annotations, with the exception of the *in vitro* p2 progenitors which mapped more strongly onto *in vivo* V2a neurons than *in vivo* p2 progenitors (**Figure 2C**). Further, the majority of our cells were identified as more mature neurons instead of progenitors, consistent with a largely post-mitotic neuronal state. Harmony integration of our RNAseq datasets clustered analogous cell types closely to one another, but did not entirely intermingle the NEP and NMP conditions (**Figure 2D**). Altogether, this analysis affirms our ability to identify regionally relevant cell types across all samples, including V2a neurons, correlate them with *in vivo* samples, and highlights the axial differences between populations derived from NEP or NMP conditions.

### Early developmental differences leave epigenetic and transcriptional marks in lineally distinct V2a neurons

Early developmental patterning events play a crucial role in molding the chromatin to position cells for subsequent stages of development. Therefore, we leveraged our snATAC-seq data to examine whether the NEP and NMP differentiation stages left long-term marks on the chromatin landscape in ways that could influence transcriptional output and neuronal function (**Figure 3A**). To focus on the V2a population, we subset all V2a cells (2672 cells) and performed differential peak analysis. This identified 1996 regions (log2FC > 1 and FDR < 0.05) that were differentially accessible between NEP and NMP V2a populations (**Figure 3B**). Examination of top differential peaks showed that peak sets included mutually exclusive regions with highly differential accessibility, not only peaks with differences in relative accessibility (**Figure 3B,C**).

**Figure 3.**
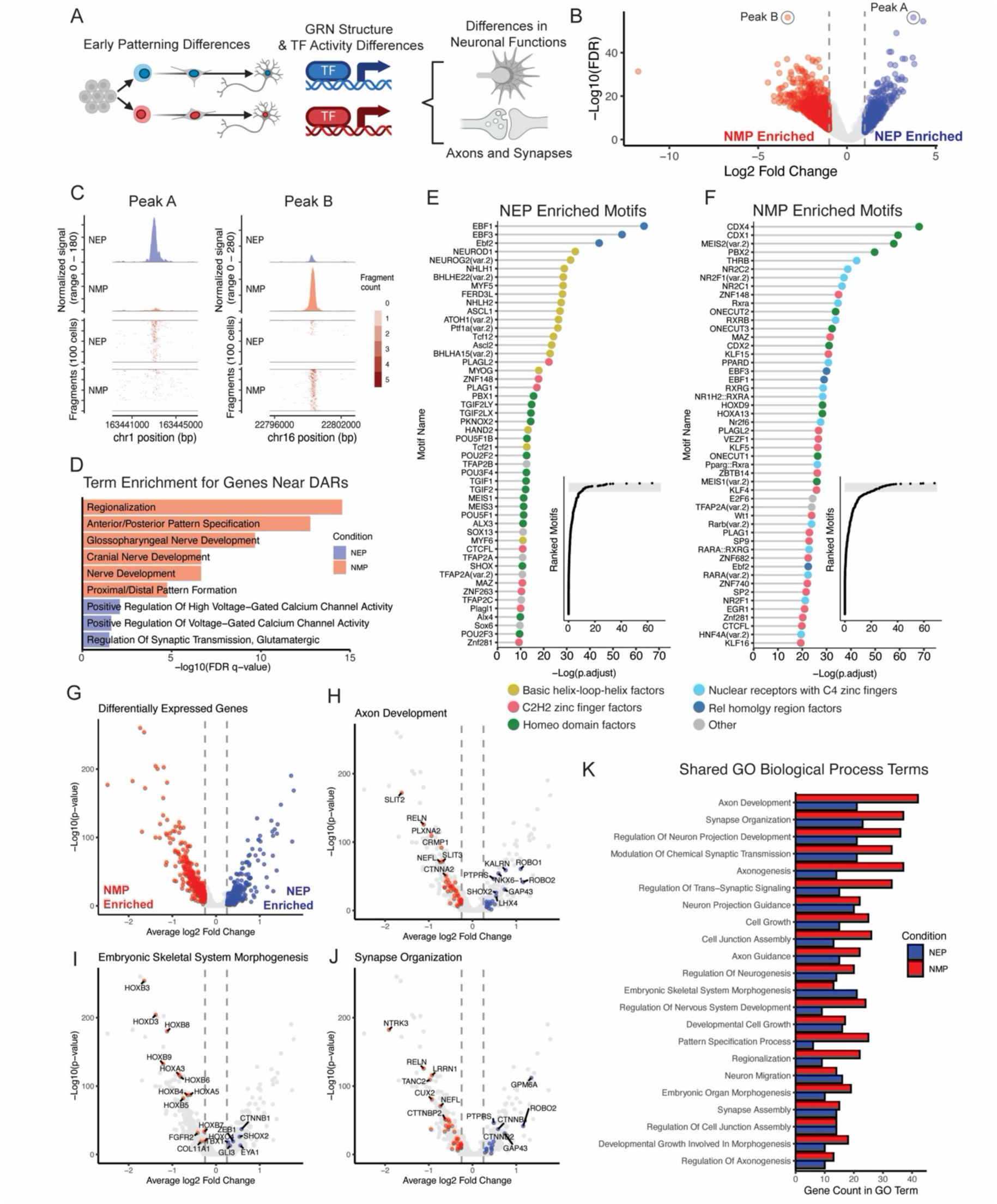
Progenitor lineage confers distinct transcription factor binding motifs and gene expression. A. Simplified schematic of the V2a differentiation and differences that could be identified using multiomic data. B. Volcano plot showing differentially accessible regions of chromatin (DAR) between NEP- and NMP-derived V2a neurons in snATAC-seq data. Blue points (positive log2 fold change) are peaks enriched in NEP samples. Red points (negative log2 fold change) are peaks enriched in NMP samples. C. Coverage plot depicting example peaks specific to either NEP or NMP conditions (left and right, respectively). Regions are represented as either normalized histograms or fragment count heatmaps per 100 cells (top and bottom, respectively). Shown peaks are also highlighted in B. D. Selected significantly upregulated gene ontology terms for NEP or NMP-specific DARs using Genomic Regions Enrichment of Annotations Tool (GREAT). E,F. Top 50 most enriched transcription factor motifs in NEP (left) or NMP-specific (right) DARs as identified in B. Each transcription factor motif is colored according to the class of transcription factor to which it belongs. Inset: ranked enrichment scores for all tested motifs, shaded gray region represents top 50 shown in the main figure. G. Volcano plot showing differentially expressed genes between NEP and NMP-derived V2a neurons in snRNA-seq data. Blue points (positive log2 fold change) are enriched in NE samples. Red points (negative log2 fold change) are enriched in NMP samples. H-J. Differentially expressed genes which belong to the indicated gene ontology (GO) term. In all cases, the GO term is significantly enriched in both NEP and NMP conditions. K. Shared significantly enriched GO Biological Process terms identified in the differentially expressed genes between NEP and NMP-derived V2a neurons. Bars represent the number of genes annotated to each respective GOBP term in each gene set.

Analysis of differentially accessible regions (DARs) did not identify differences in the relative location of peaks within genes, the distance from the nearest transcriptional start site, or the gene type annotation (protein coding, miRNA, etc.) of the nearest transcription start site (TSS) between NEP and NMP conditions (**Figure S3**). However, motif enrichment analysis of the DARs between NEP and NMP V2a neurons did identify dozens of significantly enriched transcription factor binding motifs (**Figure 3E**). In order to identify broad patterns, we annotated transcription factor (TF) motifs by class, since many related TFs bind similar motifs. In the NEP condition, motifs of Rel homology region class were the most highly enriched (EBF1/2/3), followed by basic helix-loop-helix (bHLH) class motifs, and a mix of homeodomain and zinc-finger class motifs (**Figure 3E**). In the NMP condition, homeodomain class motifs were most strongly enriched, followed by C2H2 zinc-finger and nuclear receptor classes (**Figure 3F**). The nuclear receptor class contains retinoid-related transcription factors (e.g. RXRA, RXRB, RARA) which suggests a differential action of retinoic acid in the NMP condition. While both conditions have enriched homeodomain class TFs, CDX and ONECUT family motifs are more prevalent in the NMP condition while POU and TALE family motifs are more highly enriched in the NEP condition. The bHLH family of transcription factors are known to be key regulators of neurodevelopment and several of these factors are specifically expressed during V2a development (NEUROD1, NEUROG2, ASCL1, BHLHE22; Li et al., 2005; Lu et al., 2015). However, this TF class was specifically enriched in the NEP condition (**Figure 3D**). We also tested for gene ontology enrichment using the Genomic Regions Enrichment of Annotations Tool (GREAT) to see whether nearby genes were associated with specific functions. NEP V2a neurons possessed a modest enrichment for terms such as ‘regulation of glutamatergic synaptic transmission’ and ‘regulation of voltage-gated calcium channel activity’, while NMP V2a neurons had strong enrichment for broad terms related to regionalization, anterior/posterior patterning, and nerve development (**Figure 3D, Figure S4**). These results show that the progenitor lineage of V2a neurons leaves lasting marks in the chromatin landscape even after two weeks of identical treatment. Further, these differentially accessible regions are likely to be acted upon by distinct families of transcription factors, suggesting the presence of distinct gene regulatory networks between these two conditions.

We next analyzed differences in the transcriptional networks between NEP and NMP-derived V2a neurons. Using the same set of V2a neurons, we identified 742 differentially expressed genes (**Figure 3F**; log2FC > 0.25 and Bonferroni adjusted p-value < 0.05). Consistent with our earlier findings, HOX genes were differentially expressed, reflecting distinct axial identities. Notably, differences include transcription factors, ion channels, and axon guidance molecules related to GO terms such as axon development, synapse organization, and embryonic skeletal system morphogenesis (**Figure 3H-J**). GO analysis of the differentially expressed genes identified 22 significantly enriched GO terms that are shared between the NEP and NMP conditions, yet these enrichments are driven by mutually exclusive gene sets (**Figure 3K**). These findings suggest that NEP and NMP V2a neurons use distinct gene modules to achieve similar processes, such as axonogenesis, potentially leading to different circuit architecture of neurons residing at different axial levels of the hindbrain and spinal cord. When taken with the snATAC-seq analysis, these data point to systematic chromatin changes underlying distinct transcriptional networks of V2a populations derived from NEP or NMP progenitor pools.

We next asked whether the observed differences in motif enrichment were also present in the progenitors to the V2a neurons, the p2 cells, or whether it only manifested with terminal differentiation. We therefore subset the p2 cells (781 cells) from our dataset and performed similar analyses as above. Motif enrichment in NEP p2 compared to NMP p2 DARs closely mirrored the trends observed in the V2a neurons (**Figure S5A,B**). DARs in NEP p2 cells were enriched in bHLH and homeodomain motifs, while NMP p2 regions were enriched in C2H2 zinc finger motifs and a non-overlapping set of homeodomain motifs, including CDX family genes. When we compared the DARS specific to the NEP or NMP lineage in both p2 and V2a populations, only 4.7% of NEP-specific DARs were shared between p2 and V2a neurons, while 10.7% of the NMP-specific DARs were conserved from p2 progenitors to V2a neurons (**Figure S5C,D**). Comparing the global ATAC profile of p2 and V2a populations with a Pearson correlation showed the highest degree of similarly within cell identity, not progenitor lineage (**Figure S5E)**. These data highlight that distinct progenitor lineages impart lasting differences in the regulatory network which influences chromatin accessibility to transcription factors and that these differences persist across dynamic changes in chromatin accessibility that occur over developmental time.

### Differences in population-wide synchronous activity in maturing neuron cultures

We next used calcium imaging to examine potential functional differences between NEP and NMP-derived V2a neurons. Dissociated cultures were replated at day 19 to a uniform density and allowed to form new networks for 4 days. The cell permeable dye Fluo4-AM was used to observe activity in cultures on an incubated microscope. At the day 23 timepoint, spontaneous neuronal activity was observed in both conditions, with the NMP-derived cultures eliciting a higher average number of fluorescence peaks per ROI (**Figure 4A, Movie S1, Movie S2**). However, at day 34 and day 43, the average number of peaks per ROI had equalized and became non-significant (**Figure 4A**). These results show that when ROI are assessed individually, cells in both conditions have approximately the same amount of activity as measured by peaks in Fluo4 fluorescence.

**Figure 4.**
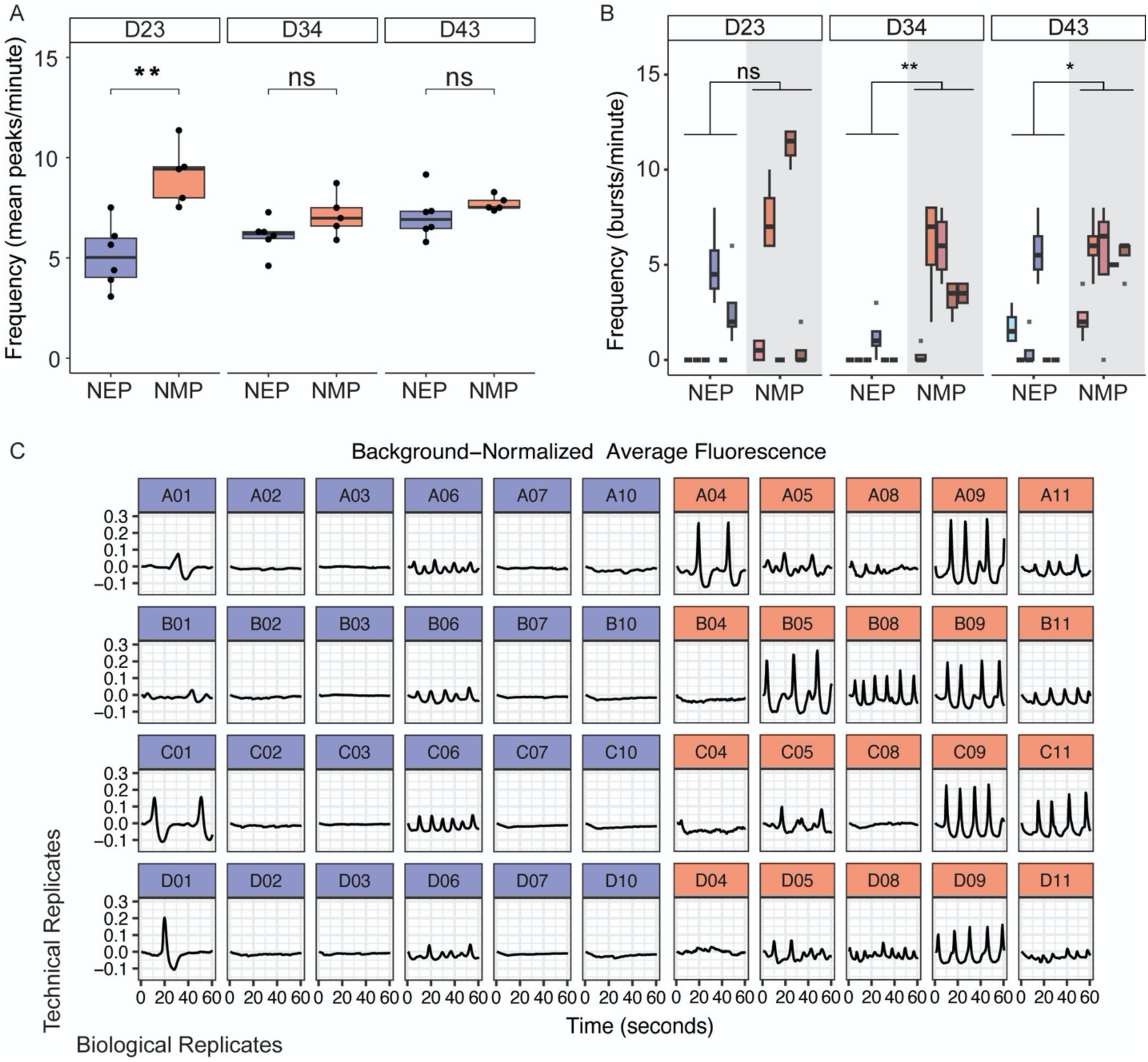
NMP-derived neurons generate robust synchronous activity in maturing cultures. A. Mean number of peaks per region of interest (ROI) captured by Fluo4-AM calcium imaging in replated V2a cultures at indicated time points. Each point represents the average of 4 technical replicates with n >= 5 biological replicates. ** is p<0.01 by Student’s t-test. B. The number of synchronous bursts per well captured by Fluo4-AM calcium imaging in replated V2a cultures at indicated time points. n >= 5 biological replicates with 4 technical replicates, each. For statistical comparison, technical replicates were collapsed to their means and compared by the Mann-Whitney U test. * is p<0.05, ** is p<0.01. C. Line graphs of background-normalized average fluorescence in each well of the data represented in Figure 4B day 43. NEP samples in blue, NMP samples in red. Columns and rows are biological and technical replicates, respectively.

However, examination of the average activity across all ROI also revealed synchronous bursts of activity across the whole culture that occurred simultaneously or in rapidly propagating waves. There was no significant difference in these bursts between NEP and NMP-derived neurons at day 23. However, we observed significantly more synchronous bursts in NMP-derived cultures at later time points compared to NEP cultures (**Figure 4B,C, Figure S6, Movie S3, Movie S4**). These observations of differential bursting activity are unlikely to be due to differences in maturation given that the NEP and NMP cultures were day-matched for differentiation and we would not expect small relative differences in maturity to exhibit such strong effects over a three-week period of continued culture. However, we do note that differentially expressed genes between the NEP and NMP V2a neurons at day 19 include genes related to synapse organization and synaptic transmission (**Figure 3K**). Also, among the most highly differentially expressed genes were subunits of cation channels and which could impact synaptic transmission (CACNA2D1 and RYR2 in the NEP condition; CACNA2D3 and KNCD2 in the NEP condition).

### Induced VSX2-expressing neurons do not resemble NEP or NMP-derived V2a neurons

Recently, the use of transcription factor overexpression to rapidly produce induced neurons has gained widespread popularity in the field (Elder et al., 2022; Fernandopulle et al., 2018; Wang et al., 2017; Zhang et al., 2013). This approach is highly efficient, scalable, and allows for the generation of neuron-like cells in just a few days, bypassing much of the complex developmental patterning typically required. Our data suggest that the developmental trajectories of the V2a precursors influence their epigenetic and transcriptional landscapes. We asked whether induced neurons that passed through minimal developmental patterning could share characteristics with either the NEP or NMP-derived populations. Based on the developmental similarities between V2a and lower motor neurons (MN), we hypothesized that removing ISL1 from the tripartite induced motor neuron cassette including NGN2, LHX3, and ISL1 could induce V2a neurons, since ectopic expression of LHX3 alone induces V2a neurons *in vivo* (**Figure 5A**) (Clovis et al., 2016; Mazzoni et al., 2013; Thaler et al., 2002).

**Figure 5.**
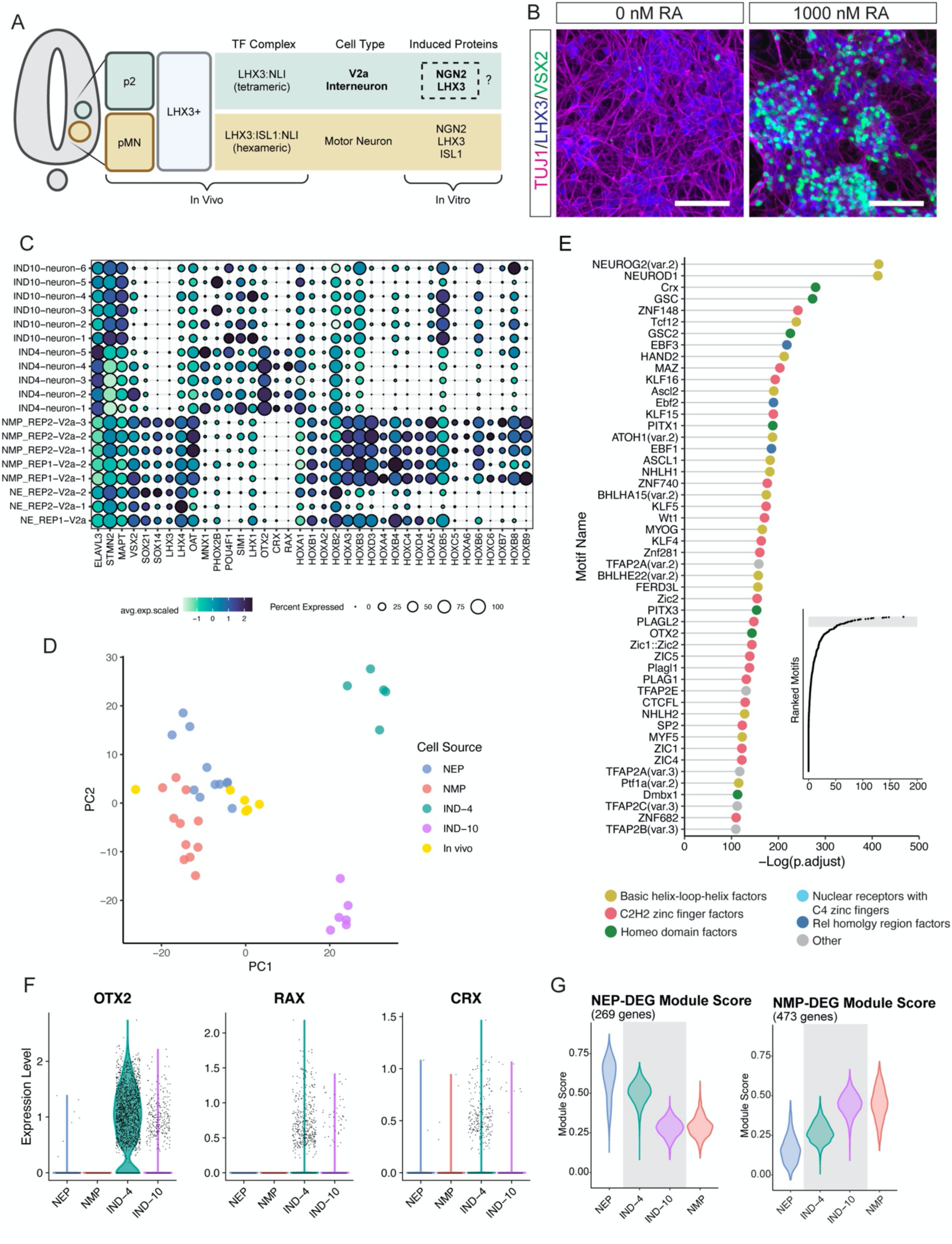
Induced neurons express anterior CNS markers and fail to recapitulate in vivo neural identities. A. Schematic of developing neural tube and the transcription factor complexes that refine V2a and motor neuron fates *in vivo*, as well as the existing and proposed transcription factor combinations to be used *in vitro*. B. Representative staining of NGN2::LHX3 induced neurons after 4 days of dox addition with either 0 nM or 1000 nM RA. LHX3 (blue) shows induced protein expression, TUJ1 (magenta) shows neural morphology, VSX2 (green) is a marker of V2a neurons. Scale bar represents 100 um. C. Comparison of NEP and NMP V2a clusters with day 4 and day 10 induced neurons. Marker genes include pan neural genes, V2a genes, off-target genes, and axial markers. D. PCA of pseudubulked clusters of NEP, NMP, *in vivo*, and day 4 and day 10 induced neurons. E. Top most significantly enriched motifs in differentially accessible regions of chromatin specific to day 4 induced neurons compared to NE and NMP V2a neurons. Each transcription factor motif is colored according to the class of transcription factor to which it belongs. Inset: ranked enrichment scores for all tested motifs, shaded gray region represents top 50 shown in the main figure. F. Expression of anterior markers OTX2, RAX, and CRX across differentiation conditions. G. Module scores of indicated populations of neurons using modules composed of the differentially expressed genes specific to either the NEP or NMP V2a populations as in Figure 3G.

Thus, we generated a plasmid containing a doxycycline-inducible NGN2::P2A::LHX3 cassette and inserted it into stem cells via piggybac transposon-mediated integration. Following 4 days of induction, we performed flow cytometry for VSX2. However, <1% of cells were positive for VSX2, despite achieving the expected neural morphology (**Figure 5B, Figure S7A**). Since PSCs default to an anterior fate, we hypothesized that our induced neurons may need an additional caudalizing signal, such as retinoic acid (RA) (Gouti et al., 2014; Metzis et al., 2018; Ozair et al., 2012). When we added graded concentrations of RA to our induced neuron differentiation protocol, we found a proportional and significant increase in the percentage of VSX2+ cells after 4 days of induction, with a maximum efficiency of ∼40% VSX2+ neurons at 1 uM RA (**Figure 5B, Figure S7A).** Notably, when we tested identical conditions on the NGN2 induced neurons and the induced motor neurons, the number of VSX2+ neurons remained <1%.

We characterized day 4 and day 10 induced V2a neurons with multiome sequencing to assess their identity. While we detected VSX2+ clusters at day 4, these cells lacked expression of other V2a markers, such as SOX21, SOX14, and endogenous LHX3 (**Figure 5C, Figure S7B,C**). The VSX2-clusters were largely positive for the motor neuron marker MNX1 (**Figure 5C, FigureS7B,C**). While this was not the target population, the shared LHX3+ lineage of both cell types *in vivo* supports the idea that there could be a balance between these populations produced by overexpression of LHX3 in stem cells. More unexpected was the observation of PHOX2B and POU4F1 overlapping VSX2 and MNX1 populations, respectively, as well as the presence of other interneuron markers SIM1 and LHX1 (**Figure 5C, Figure S7C,E**). Expression of PHOX2B and POU4F1 have been described in studies of NGN2-induced neurons and may be a product of the induced method (Lin et al. 2021). It is unclear if SIM1 and LHX1 are expressed under the same conditions. Nevertheless, these markers persist through day 10 and overlap remaining VSX2+ and MNX1+ clusters. To define the global similarities between induced and directed NEP and NMP neurons generated *in vitro*, as well as to *in vivo* spinal neurons, we psuedobulked the RNA data from neuron clusters and embedded them in PCA space (**Figure 5D**). The PCA showed that both of the induced time points clustered distantly from the NEP, NMP, and *in vivo* clusters along PC1, as well as distantly from each other along PC2. By contrast, both NEP and NMP samples were close to *in vivo* clusters.

The motifs enriched in the differentially accessible peaks between day 4 induced neurons and combined NEP and NMP V2a neurons showed an enrichment of anterior homeodomain motifs such as GSC, CRX, and OTX2 in addition to the expected bHLH motifs from NGN2 overexpression (**Figure 5E**). At day 4, we noticed significant expression of the anterior gene OTX2 along with lower levels of eye-related factors CRX and RAX while HOX gene expression mostly stopped at HOXB2 (**Figure 5C,F**). However, day 10 induced neurons downregulated OTX2, CRX, and RAX and increased expression of HOXB5 and HOXB8, suggesting that they adopted a more caudal identity following an additional 6 days in culture in the presence of retinoic acid (**Figure 5C,F**). This caudal shift in identity from day 4 to day 10 was further supported by module scoring using the significantly differentially expressed genes between the NEP and NMP V2a neurons (**Figure 5G**).

These data show that induced neurons possess a superficial similarity to V2a neurons. Upon close inspection, these cells lack co-expression of secondary marker genes (SOX14 and SOX21) and express transcription factors that mark other cell populations (PHOX2B and POU4F1; **Figure 5C**). Further, the axial identity of these neurons appears to be labile after achieving a morphologically ‘neuron-like’ post-mitotic state (**Figure 5G**). This contrasts with normal developmental progression, where neural progenitors fail to acquire more caudal identities after an early signaling window (Metzis et al., 2018). While we did not exhaustively test transcription factor and small molecule combinations that could produce more robust V2a-like neurons, these experiments highlight that the most common V2a marker, VSX2, induced by LHX3 in a developmentally relevant progression is not sufficient to establish an underlying gene regulatory network that resembles that of a V2a neuron and that induced neurons are still susceptible to external cues. These findings prompted us to identify additional factors that govern the acquisition of lineally distinct V2a identity.

### In silico GRN modeling predicts novel regulators of lineally distinct V2a neuron signatures

Having observed that different progenitor lineages conferred hundreds of differentially expressed genes and access to context-specific transcription factor motifs in post-mitotic neurons, we next sought to identify the gene regulatory network (GRN) responsible for these differences. For this, we turned to CellOracle, a computational tool which takes snRNA-seq and snATAC-seq data to construct population-specific GRNs based on correlating RNA expression and chromatin peak co-accessibility (Kamimoto et al., 2023). After building GRNs, individual TFs can be ‘knocked out’ *in silico* in order to predict changes in cell state or differentiation trajectory (**Figure 6A**).

**Figure 6.**
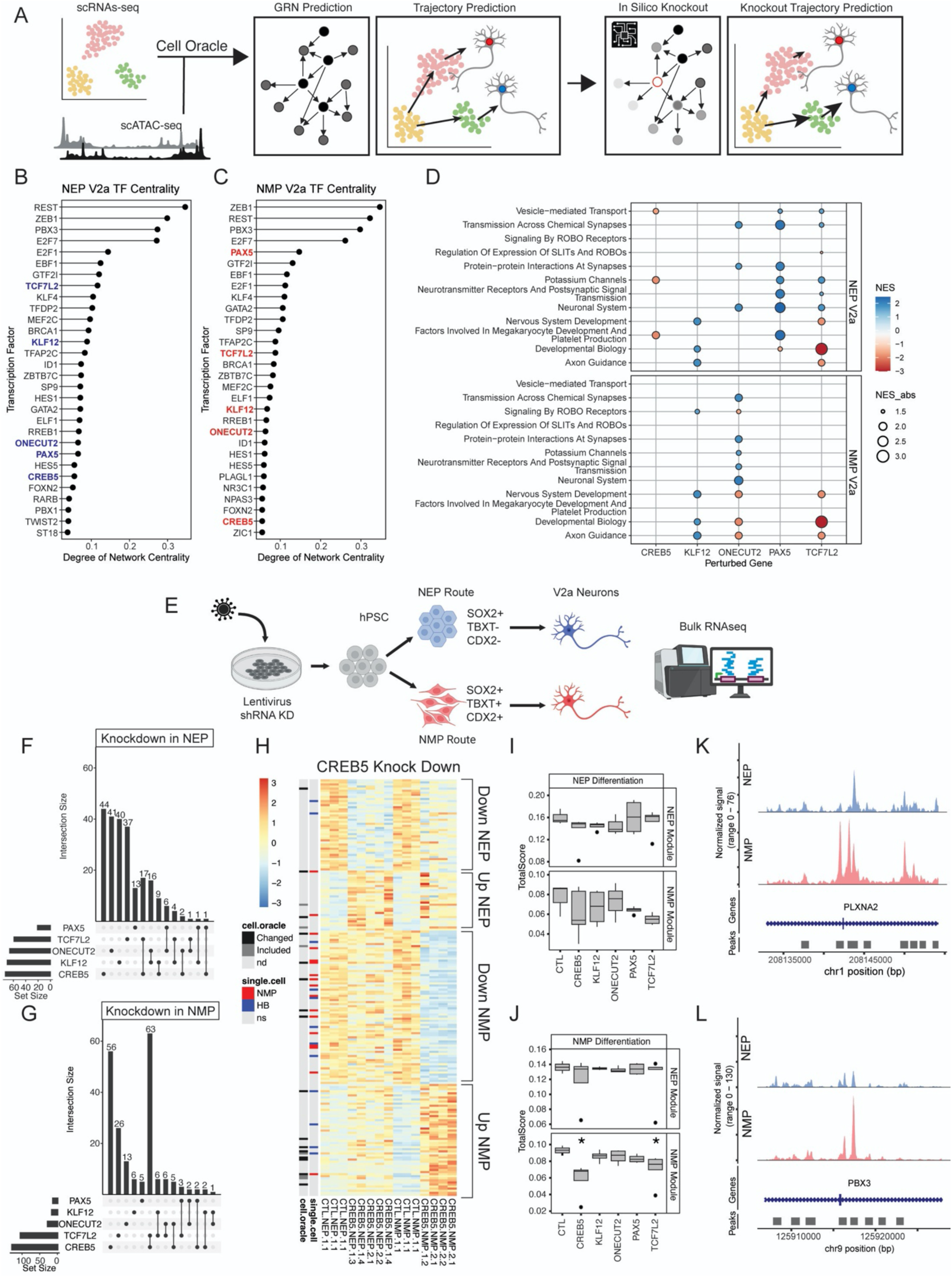
CREB5 and TCF7L2 regulate an NMP-enriched V2a gene signature. A. Schematic of the CellOracle pipeline. Single nucleus RNA- and ATAC-seq datasets are incorporated to generate predicted gene regulatory network (GRN) and map default trajectories. Specific nodes of the GRN can then be computationally ‘knocked-out’ and the effects propagated throughout the regulatory network. These effects can also be visualized as differences in the vectorized trajectory of cell populations. B,C. Top 30 transcription factors ranked by network centrality score in the predicted GRNs for both NEP and NMP V2a neurons, respectively. Highlighted genes were selected for further validation. D. Gene set enrichment analysis (GSEA) in NEP and NMP V2a neurons after in silico gene perturbation (knockout). Shown are the normalized enrichment scores (NES) for selected gene sets. E. Schematic showing validation experiments using lentiviral knockdown of selected targets. F. Upset plot depicting the number and overlap of differentially expressed genes between control and knockdown of the indicated gene under NEP differentiation conditions. G. Upset plot depicting the number and overlap of differentially expressed genes between control and knockdown of the indicated gene under NMP differentiation conditions. H. Heatmap depicting the significantly differentially expressed genes in the CREB5 KD condition across both NEP and NMP differentiations. I. Boxplots of single sample GSEA scored only on NEP condition knockdown samples using the differentially expressed genes between NEP and NMP V2a neurons as indicated modules. No sample is significantly different from the control condition by Student’s t-test. J. Boxplots of single sample GSEA scored only on NMP condition knockdown samples using the differentially expressed genes between NEP and NMP V2a neurons as indicated modules. The NE module is not significantly changed in any knockdown sample, but both CREB5 and TCF7L2 knockdown samples score significantly lower in the NMP module than controls. * is p < 0.05 by Student’s t-test. K,L. Coverage plots centered on differentially accessible regions enriched in the NMP condition which are located within genes (PLXNA2 and PBX3, respectively) which are significantly downregulated with NMP CREB5 KD samples.

Focusing on the V2a populations, we used CellOracle to produce separate GRNs for the NEP and NMP V2a populations using the top 3000 most variably expressed genes. We then examined which transcription factors contained the highest centrality scores in the predicted network, with higher scores correlating with larger numbers of total regulatory connections (**Figure 6B,C**). Both of the V2a populations contained largely overlapping lists of highly central TFs, likely a result of their shared identity. To investigate which of these TFs may explain the observed condition-specific differentially expressed genes, we knocked down each of the top 25 most central genes in NEP or NMP V2a populations *in silico*. We then performed Gene Set Enrichment Analysis (GSEA) on the perturbed conditions to identify what biological processes were up or down-regulated upon knockout (**Figure 6D, Figure S8**). Several GSEA terms related to neural function were enriched in either the NEP or NMP condition by single gene knockout (**Figure 6D, Figure S8**). In order to experimentally validate these computational predictions, we narrowed down the list of potential targets using several criteria including expression level, whether or not the gene was differentially expressed in the RNA-seq data, whether the TF or similar TF motifs were enriched in the ATAC-seq data, and centrality in both conditions. We selected PAX5, CREB5, KLF12, ONECUT2, and TCF7L2 for further validation (highlighted in **Figure 6B-D**). We knocked down these factors using short hairpin RNAs (shRNA) integrated into hPSCs using lentiviral vectors. hPSCs were then differentiated along the NEP or NMP routs toward V2a neurons (**Figure 6E**) and bulk RNA was collected for sequencing on day 19.

Next, we examined the genes that were differentially expressed in NEP or NMP knockdown conditions (**Figure 6F,G, Figure S9**). Contrasting with the NEP condition where the majority of differentially expressed genes (DEG) were specific to knockdown of one gene (**Figure 6F**), we noticed that knockdown of either CREB5 or TCF7L2 in the NMP condition resulted in the largest total numbers of DEG, many of which were overlapping (**Figure 6G**). A heatmap of the DEG in the NEP and NMP CREB5 knockdown conditions showed that the NEP-specific DEG exhibited similar trends in the NMP conditions, although not statistically significant (**Figure 6H**). The NMP-specific DEG, however, had highly specific effects, and were also enriched for the DEG identified in our multiome data (**Figure 6H**). Similar observations were made for the TCF7L2 knockdown (**Figure S9A**). While neither the NEP or NMP module was significantly altered in the NEP differentiations, both CREB5 and TCF7L2 knockdowns significantly reduced the NMP module in NMP differentiations (**Figure 6I,J**). Thus, it appears that CREB5 and TCF7L2 play key regulatory roles specific to the NMP lineage and that they contain largely overlapping regulons.

Lastly, we examined the list of differentially accessible regions between NEP and NMP V2a neurons and filtered it for peaks that were annotated in or nearest to the DEG in the NMP CREB5 knockdown samples. This identified 25 NMP-enriched peaks within or near 15 unique genes downregulated by CREB5 KD, including PLXNA2, a semaphorin receptor which plays a role in axon guidance, and PBX3, a transcription factor which can form complexes with HOX genes and is thought to play a wide role in early development (**Figure 6K,L**). These findings suggest that CREB5 and TCF7L2 could regulate some of their DEG by either opening unique regions of chromatin or by binding to these peaks to regulate transcription.

## Discussion

Our study demonstrates that distinct developmental trajectories in the differentiation of V2a neurons result in lasting, lineage-specific transcriptional and functional differences. Using two distinct progenitor differentiation methods—one generating NEP and the other NMP—we observed that these lineages exhibit differential gene expression, chromatin accessibility, and network activity, even under identical culture conditions during the later stages of differentiation.

Our multiomic analysis revealed that early lineage differences imprint unique chromatin landscapes that persist into the mature neuron stage, suggesting that early developmental events impart structural changes in chromatin that influence long-term gene regulatory networks. The NMP-derived V2a neurons, for example, displayed enrichment in transcription factor motifs associated with homeodomain-containing transcription factors, which are critical for regionalization and spinal cord identity. Conversely, NEP-derived neurons showed a broader distribution of bHLH and Rel homology motifs, consistent with a hindbrain-like identity. This discovery underscores the impact of progenitor origin on the epigenetic landscape, potentially guiding functional differentiation even within seemingly similar neuronal subtypes.

Functional assessments showed that NMP-derived V2a neurons displayed more rapid establishment of synchronous network activity compared to their NEP counterparts. This increased network activity may reflect differences in axonal connectivity or intrinsic excitability, which could influence their suitability in regenerative therapies targeting motor or sensory circuits. These functional differences highlight the importance of considering progenitor origin in therapeutic applications, where specific activity profiles may be desirable.

Future functional studies should explore *in vivo* validation of NEP and NMP-derived neurons to assess circuit integration and functional synaptic integration post-transplantation. Additionally, our computational analysis identified CREB5 and TCF7L2 as potential regulators of NMP-specific gene networks in V2a neurons, warranting further investigation into their role in modulating neurodevelopmental pathways. These insights may aid in the development of improved differentiation protocols for generating functionally specialized V2a neurons.

Our experiments with induced VSX2-expressing neurons, which bypassed key developmental patterning stages, yielded cells that displayed some superficial characteristics of V2a neurons but differed significantly in transcriptional identity. The axial identity of these induced neurons remained labile, indicating that essential developmental cues are necessary to establish and maintain regional specificity. These findings are consistent with previous studies showing that induced neurons often lack the stable chromatin modifications characteristic of lineage-specific differentiation. Our results suggest that induced neurons may not fully replicate the gene regulatory networks of authentic V2a neurons, raising concerns about their applicability in disease modeling and therapeutic discovery where functional specificity is critical. Future studies using *in silico* GRN modeling and perturbation could identify additional transcription factors that can be overexpressed to enhance the fidelity of induced V2a neurons. By pinpointing key regulators that drive lineage-specific traits, this approach could provide insights into constructing more stable and functionally accurate V2a neuron populations, ultimately improving their utility in disease models and therapeutic applications.

In summary, our study establishes a foundational framework that extends beyond the specific context of V2a neurons, underscoring the importance of refining hPSC differentiation strategies. Our findings emphasize the necessity of generating cells that not only express key lineage markers but also faithfully replicate the developmental stages and processes encountered during fetal development. Enhancing the specificity and accuracy of differentiations will enable more precise and diverse basic and translational applications.

### Limitations of the study

Our study design enables us to isolate the effects of NEP and NMP progenitor lineages on V2a neurons; however, our multiomic analysis captures only a single time point. While we observe population heterogeneity and can separate p2 populations for pseudotemporal analysis, we cannot conclusively determine whether lineage identity is retained over time. snATAC-seq reveals enriched motifs in specific populations, but experiments such as ChIP-seq are needed to pinpoint TF binding at specific loci. Although we observed distinct activity patterns between maturing NEP and NMP cultures, these findings are confounded by mixed populations. Variations in motor and interneuron proportions may influence burst activity, necessitating targeted drug perturbations or sorted cultures to resolve these effects.

## Author Contributions

N.H.E. designed, completed, and analyzed all *in vitro* experiments including cell line editing, differentiation, 2D staining, flow cytometry, calcium imaging, and RNA extraction. Designed and executed Mutliome data analysis, bulk RNA sequencing analysis, and calcium imaging analysis. Generated all figures and wrote the manuscript. A.M. performed all in silico analysis using the CellOracle tool and assisted in figure design. E.A.B. designed and assisted in the creation and validation of the FONX4:Cre reporter cell line. R.M.S assisted with lentiviral production and provided scripts for Singscore analysis. Assisted with figure generation and design. L.V.Z. advised on experimental design and data interpretation. T.C.M. designed and conceived the study and oversaw the generation of the FOXN4:Cre reporter cell line and differentiation optimization experiments. F.F. designed and conceived the study, oversaw experiments, and wrote the manuscript.

## Acknowledgements

We would like to thank Kenneth Wu and Luke Judge for generating and sharing the XR1 cell line. We would like to thank Karissa Hansen for her expertise and assistance with bulk RNAseq analysis. We also thank following core facilities for their support and experimental advice: the UCSF Center for Advanced Technology (CAT) supported by UCSF PBBR, RRP IMIA, and NIH 1S10OD028511-01 grants, UCSF Laboratory for Cell Analysis, UCSF Center for Advanced Light Microscopy, UCSF Genomics CoLabs, UCSF Institute for Human Genetics, Gladstone Stem Cell Core, Gladstone Assay Development Core, and Gladstone Flow Core. We would like to thank the members of the Fattahi Lab for their thoughtful feedback. The work was generously supported by grants from the NIH Director’s New Innovator Award (DP2NS116769) and the National Institute of Diabetes and Digestive and Kidney Diseases (R01DK121169) to F.F. N.H.E was supported by a fellowship from the ARCS Foundation. L.V.Z was supported by a grant from the California Institute for Regenerative Medicine (DISC2-14180).

---

**Supplemental Figure 1.**
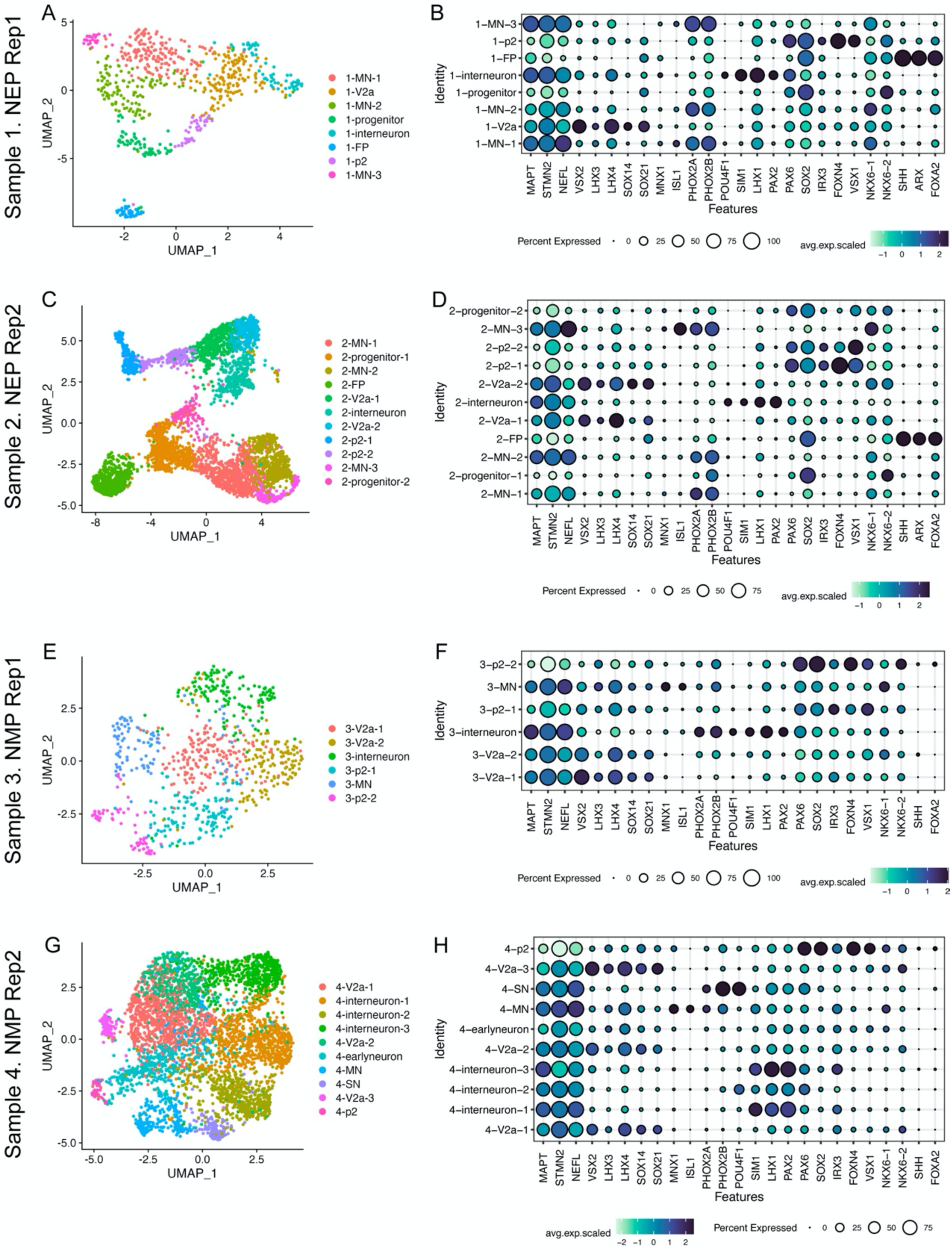
Individual sample clustering and cell annotation by marker gene sets. A, C, E, G. UMAP plots of the indicated sample with clusters identified using resolution = 0.8. B, D, F, H. Dot plots showing the expression of marker gene sets used to classify cell types within the indicated sample.

**Supplemental Figure 2.**
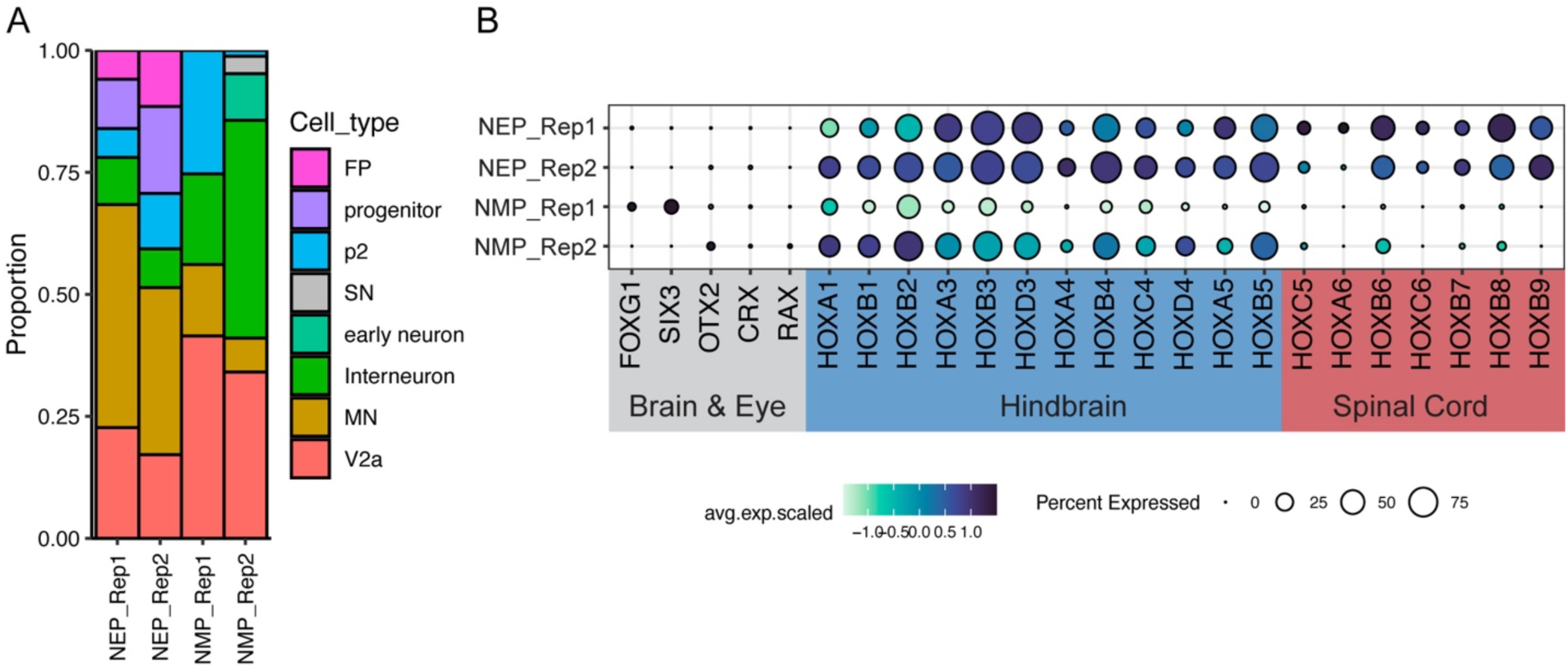
Characterization of population proportion and axial identity in multiome samples. A. Expression of anterior-posterior axial markers in each multiome sample. Colored groups of genes indicate approximate anatomical boundaries. B. Bar plot showing the proportion of each annotated cell type for each multiome sample.

**Supplemental Figure 3.**
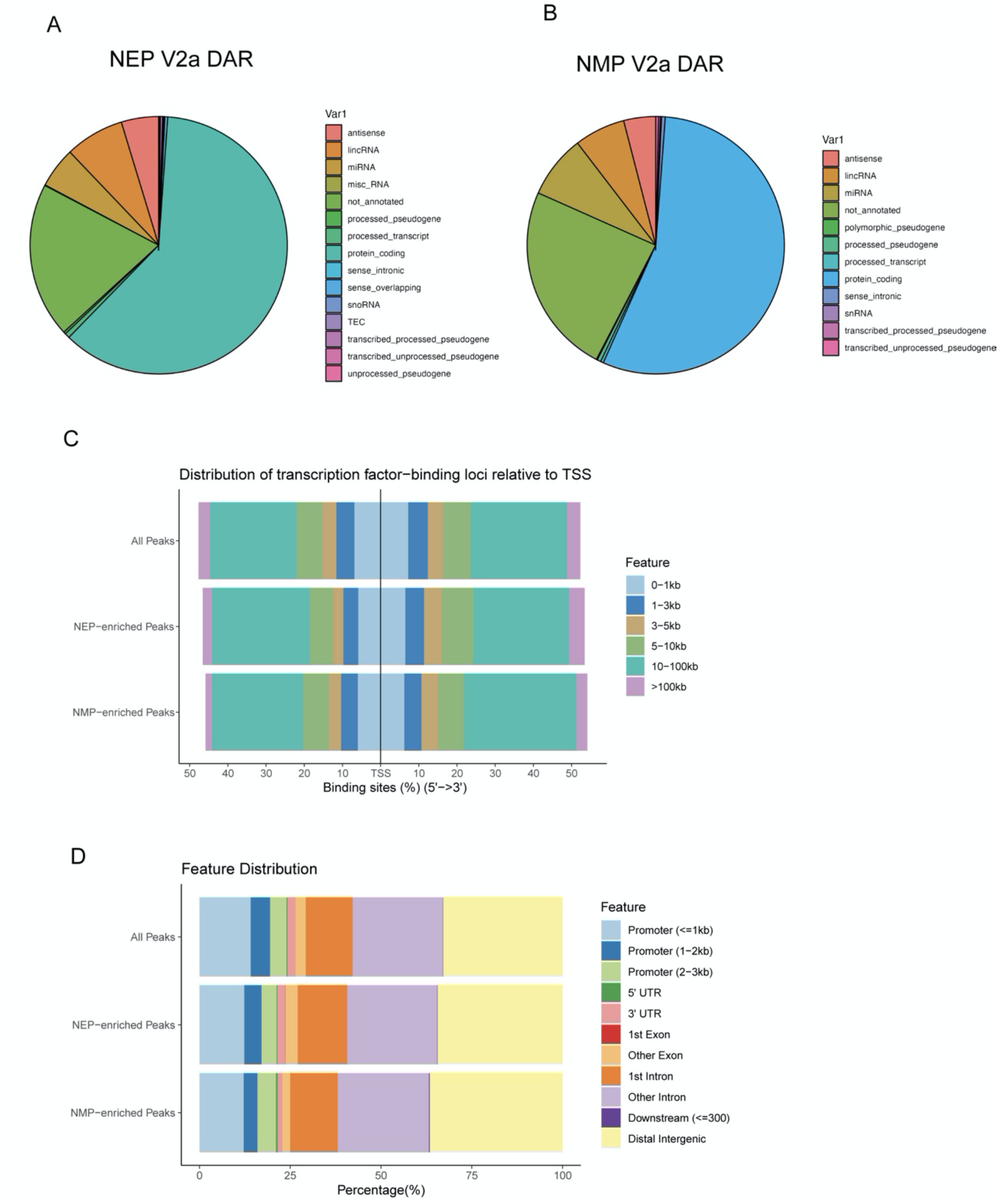
Similarities in the annotation and relative location of differentially accessible ATAC peaks between NEP and NMP V2a neurons. A, B. Pie chart of the gene type annotation for the gene with the nearest TSS to differentially accessible chromatin peaks between NEP and NMP-derived V2a neurons. C. Bar plot showing the distribution of differentially accessible peaks between NEP and NMP V2a neurons. D. Bar plot showing the relative location of differentially accessible peaks within genes between NEP and NMP V2a neurons

**Supplemental Figure 4.**
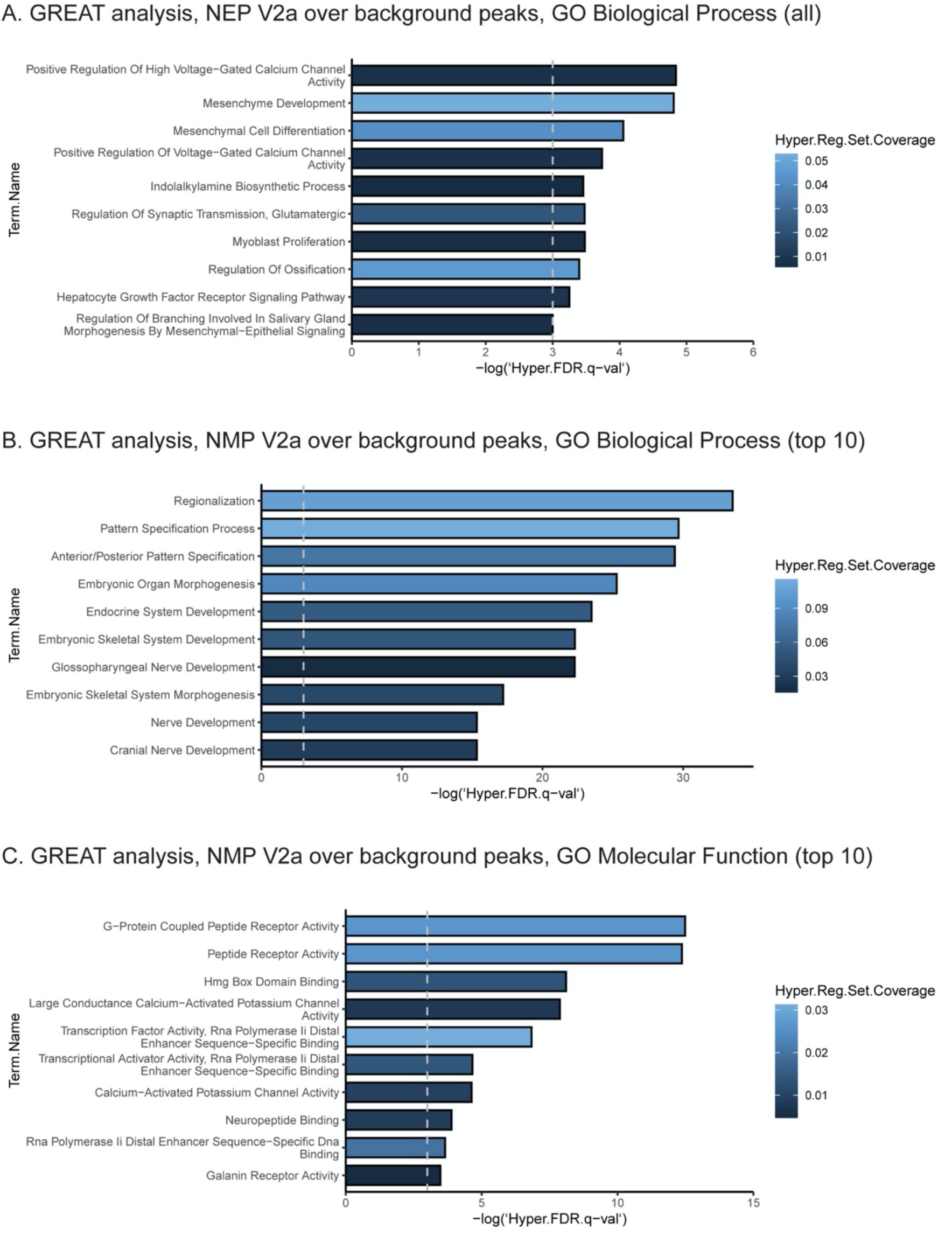
Enrichment of Gene Ontology terms for the genes near ATAC peaks. A. GREAT Analysis showing all significantly enriched biological process GO terms for the genes near NEP-enriched peaks. Dashed grey line is significance threshold. B. GREAT analysis showing the top 10 significantly enriched biological process GO terms for the genes near NMP-enriched peaks. Dashed grey line is significance threshold. C. GREAT analysis showing the top 10 significantly enriched molecular function GO terms for the genes near NMP-enriched peaks. Dashed grey line is significance threshold.

**Supplemental Figure 5.**
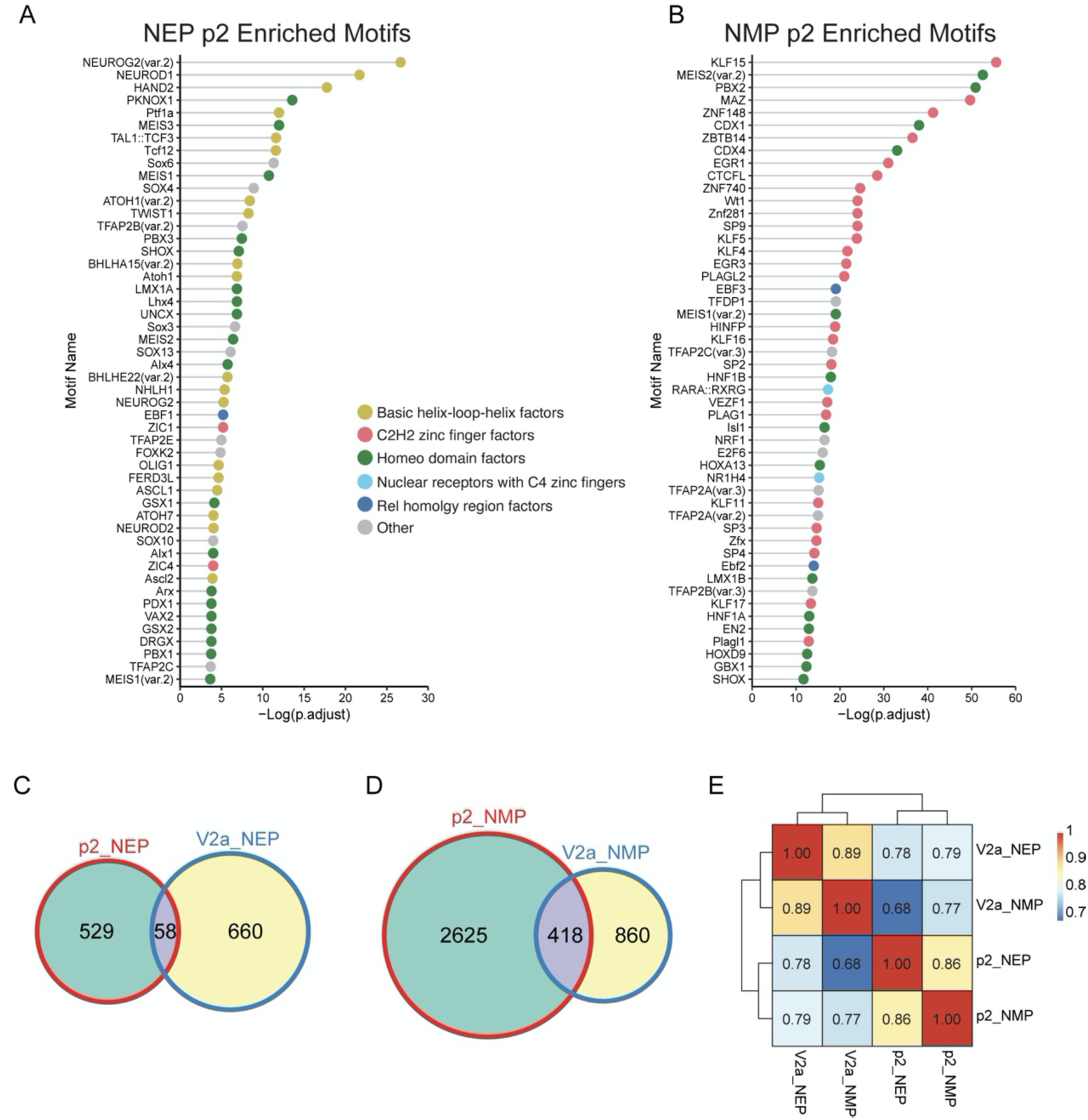
Lineally distinct p2 progenitors possess enriched transcription factor motifs similar to V2a but limited overlap in lineage specific ATAC peaks. A. Top 50 most enriched transcription factor motifs in NEP p2 DARs. Each motif is colored by transcription factor class. B. Top 50 most enriched transcription factor motifs in NMP p2 DARs. Each motif is colored by transcription factor class. C. Venn diagram of the DARs found in both the p2 and V2a cells from the NEP condition D. Venn diagram of the DARs found in both the p2 and V2a cells from the NMP condition E. Pearson diagram showing the global similarity of ATAC peaks between p2 and V2a populations.

**Supplemental Figure 6.**
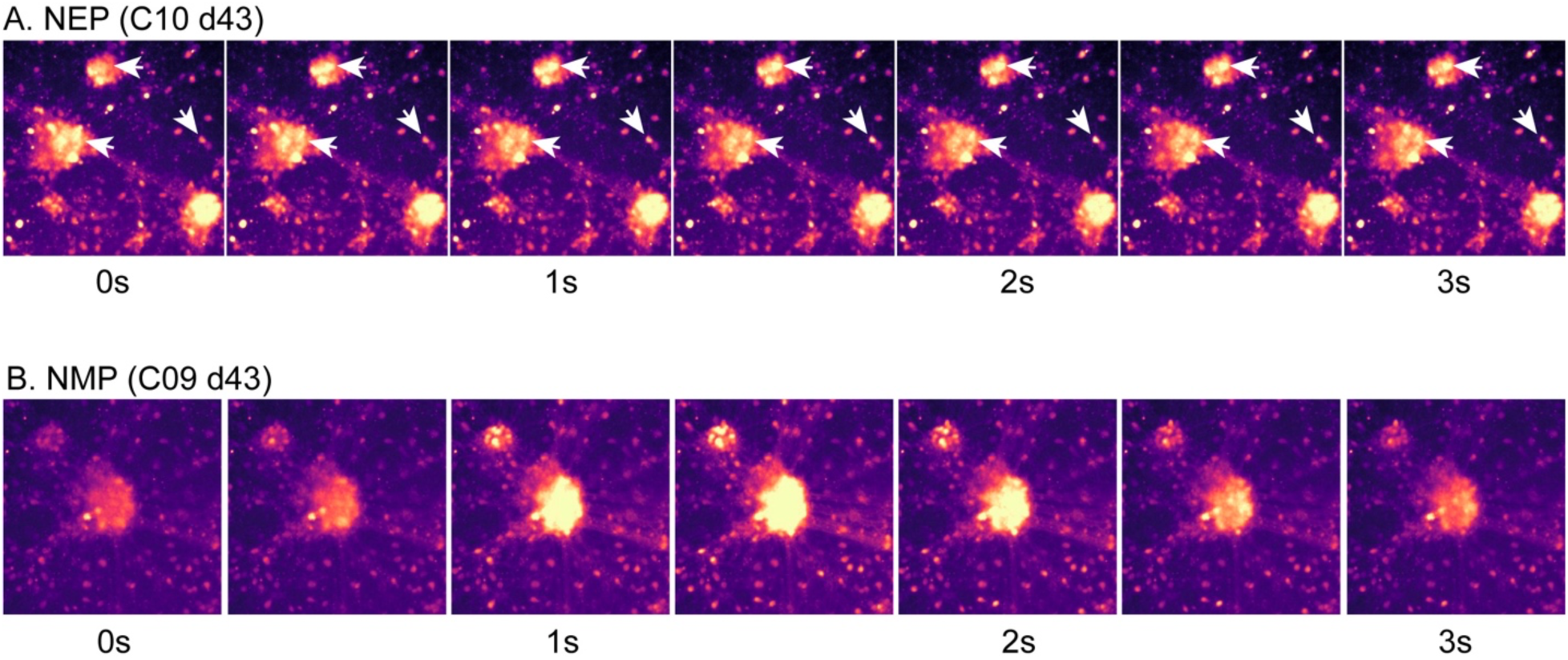
Representative stills of neural activity from calcium imaging videos. A. Representative images from a d43 NEP-derived neuron culture calcium imaging video. Arrow heads point to regions with fluorescent activity. B. Representative images from a d43 NMP-derived neuron culture calcium imaging video. Panels show a synchronous burst of activity across the population.

**Supplemental Figure 7.**
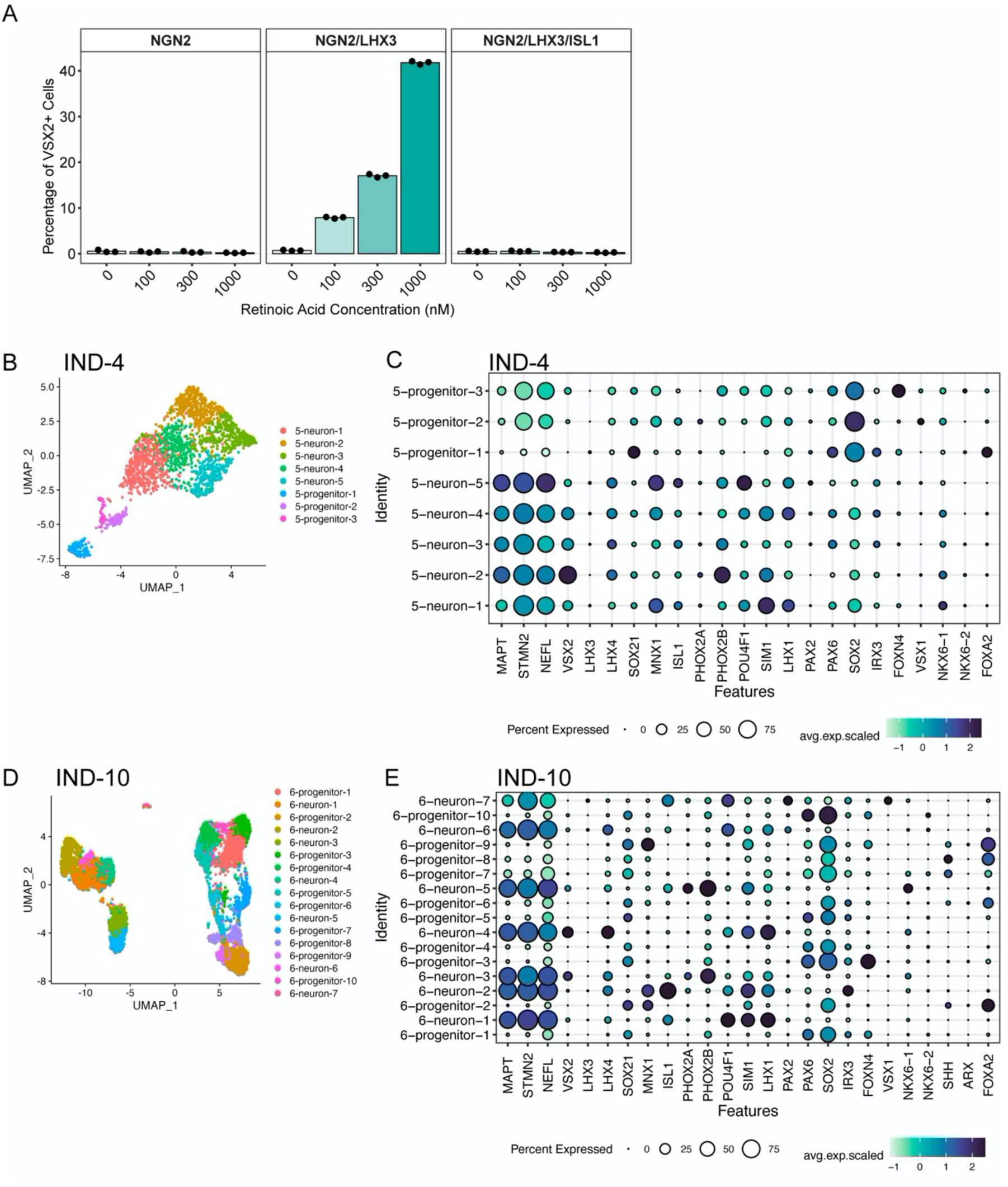
Marker gene expression in induced neurons. A. Flow cytometry quantification of VSX2 expression in induced neuron populations under graded concentration of retinoic acid (RA). The specific combination of transcription factors is shown at the top of each panel. n = 3 technical replicates. B. UMAP of day 4 induced neurons. C. Marker gene expression in clusters identified in day 4 induced neurons D. UMAP of day 10 induced neurons. E. Marker gene expression in clusters identified in day 10 induced neurons.

**Supplemental Figure 8.**
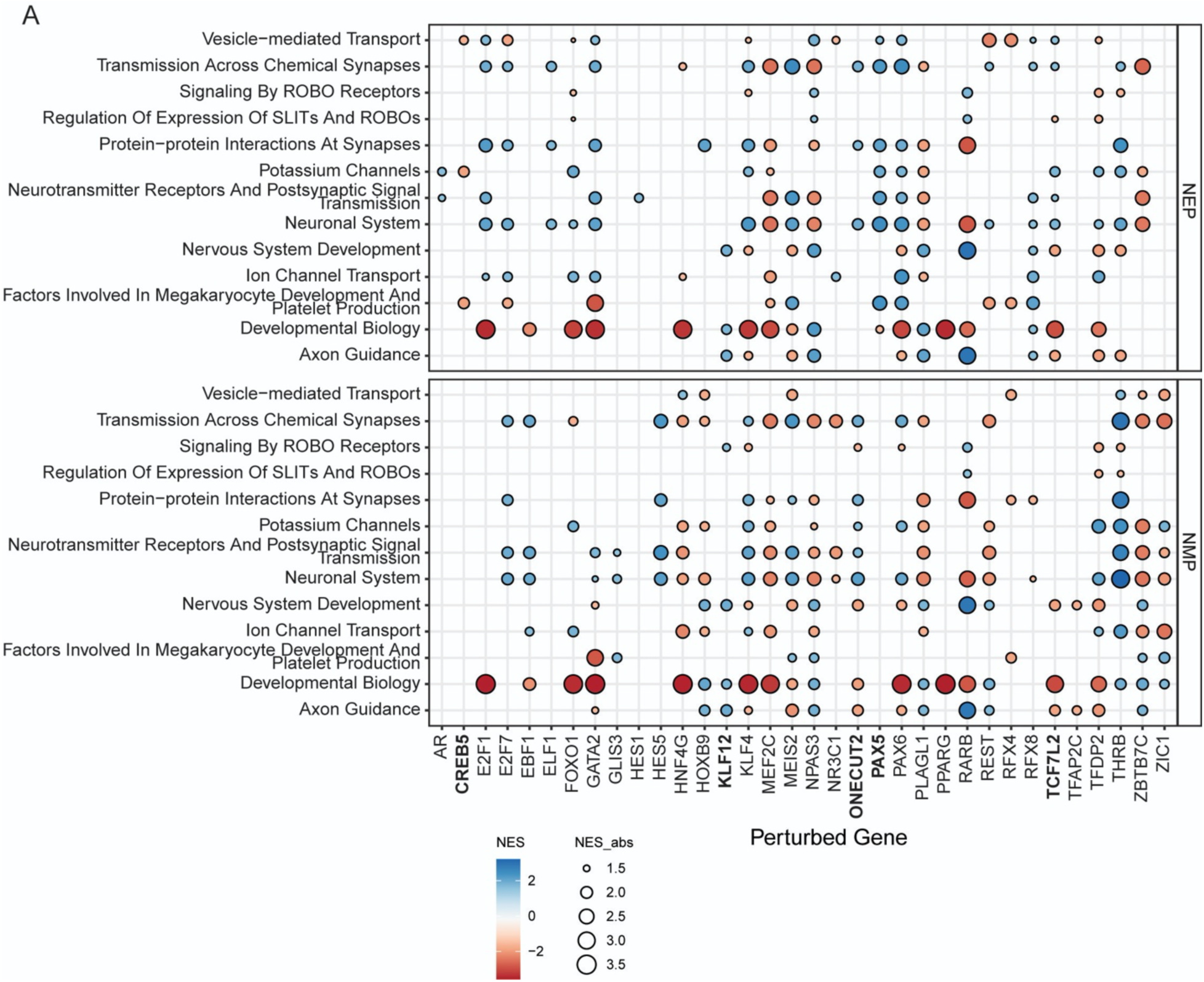
GSEA in distinct V2a populations after in silico knockout of highly central transcription factors. A. GSEA normalized enrichment scores for selected gene sets after predicting CellOracle gene knockout effects in both NEP and NMP V2a neurons. Target genes for CellOracle knockout are from the most central genes in the predicted V2a gene regulatory networks.

**Supplemental Figure 9.**
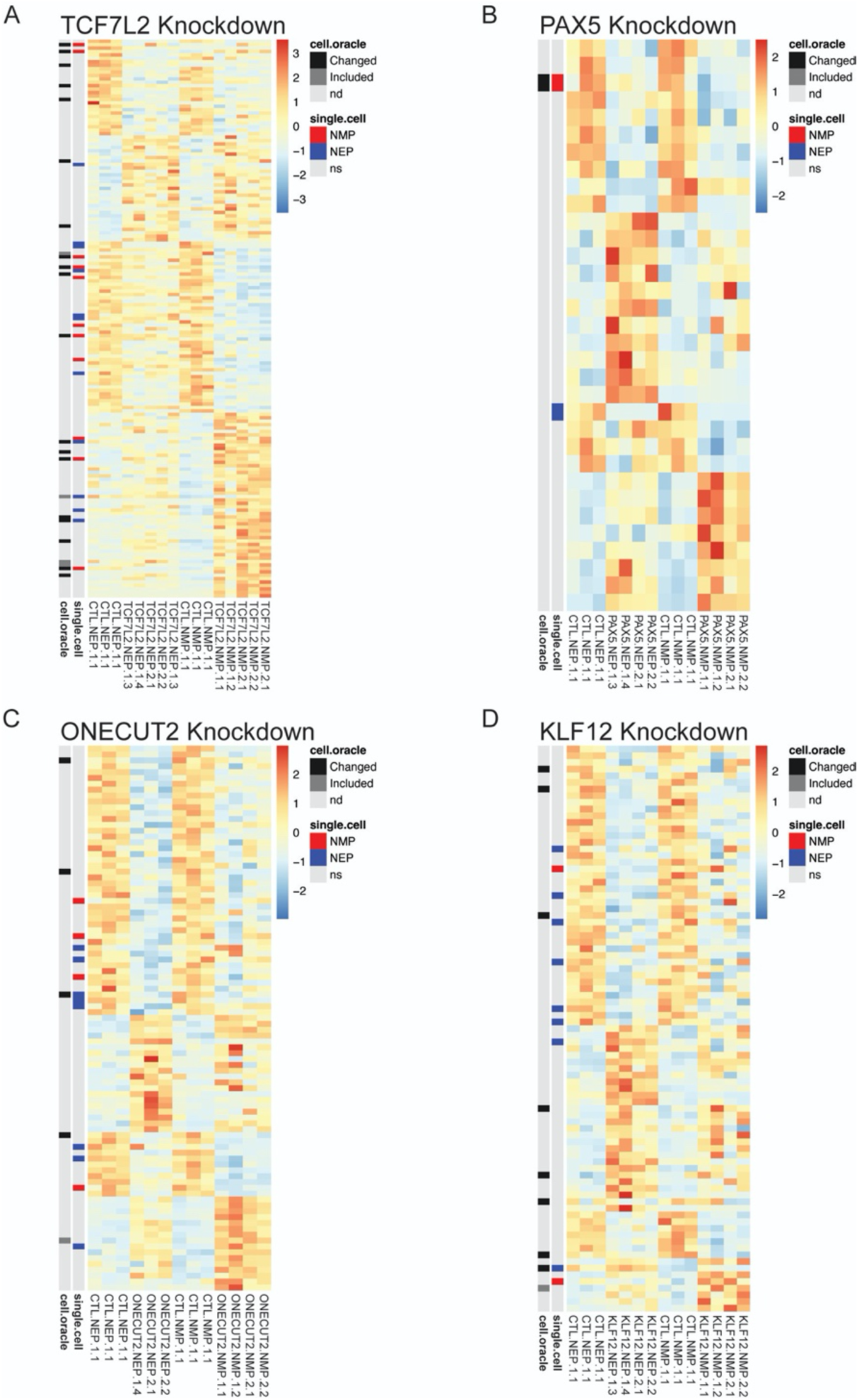
Differential gene expression resulting from transcription factor knockdown in NEP and NMP differentiations. A. Heatmap depicting the significantly differentially expressed genes in the TCF7L2 KD condition across both NEP and NMP differentiations. B. Heatmap depicting the significantly differentially expressed genes in the PAX5 KD condition across both NEP and NMP differentiations. C. Heatmap depicting the significantly differentially expressed genes in the ONECUT2 KD condition across both NEP and NMP differentiations. D. Heatmap depicting the significantly differentially expressed genes in the KLF12 KD condition across both NEP and NMP differentiations.

## Methods

### Stem cell maintenance

The parental stem cell lines used in this study include WA09 (WiCell), WTC11 (Coriell Institute), and WTB lines (Conklin Lab; <u>Miyaoka et al., 2014)</u>. Stem cells were seeded on Geltrex coated plates and maintained in mTeSR1 with daily feeds or in mTeSR Plus and fed every other day and kept at 37°C and 5% CO2. Stem cells were passaged at ∼70% confluency using Accutase at a 1:10 ratio. A defined pro-survival small molecule cocktail, CET, was included in the media for the first 24 hours after passaging <u>(Chen et al., 2021)</u>. All stem cell lines were tested quarterly for mycoplasma.

### Generation of the FOXN4:GFP reporter line

The FOXN4:Cre cell line was built into the WTC-hNIL transgenic iPSC line (previously described in Shi et al., 2018; Fernandopulle et al., 2018). First, a lox-STOP-lox reporter cassette called XR1(Addgene plasmid #129111; a gift from Bruce Conklin) was integrated into the CLYBL safe harbor locus using locus-specific TALEN (Addgene plasmid #62196 and #62197; gifts from Jizhong Zou). Donor and TALEN plasmids were nucleofected into WTC-hNIL stem cells using the P3 Primary Cell 4D Nucleofector X Kit (Lonza). Cells were then selected for with G418 and integration was confirmed using junction PCR (**Supplemental Table 3**). Excision of the lox-STOP-lox cassette was tested using CRISPR-mediated activation of the reporter by excision of the terminator using combinations of 5 RNA guides targeting each side of the cassette (L1-5, R1-5; **Supplemental Table 3**). We next made a donor plasmid for integration of a P2A-cleaved Cre in-frame with the endogenous FOXN4 locus. The donor plasmid contained a pCAGG::mCherry integration reporter for non-antibiotic selection which could be removed by CRISPR/Cas9 using flanking tia1L guide sites (Roberts et al., 2019, fragments taken from AICSDP-45: TTN-mEGFP, a gift from Allen Institute for Cell Science; Addgene # 114412). A homology region of ∼2kb in length surrounding the FOXN4 stop codon was amplified using PCR for HDR targeting (**Supplemental Table 4**). FOXN4:Cre donor plasmid was nucleofected into XR1-containing stem cells along with Cas9 complexed with a single-guide RNA (ACAGCUCAAAGCAGGGCUAU; Synthego). After recovery and expansion, mCherry+ clones were hand-picked and confirmed with junction PCR. To remove the mCherry selection cassette, tia1L sgRNA (GGUAUGUCGGGAACCUCUCC, Synthego) and Cas9 were added using Lipofectamine Stem to mCherry+ cultures. After one passage, mCherry-cells (∼30% of the population) were sorted on a FACS Aria Fusion, recovered, expanded, and cryopreserved.

### Neuromesodermal progenitor differentiation

Neuromesodermal progenitor (NMP) differentiation loosely followed the protocol set out in Lippmann et al. (2015). Briefly, stem cells were passaged as normal in 6-well plates and allowed to grow to ∼70% confluency in mTeSR Plus. To begin differentiation, cells were then fed with 4 mL/well E6 media. One day later, cells were fed with 4 mL/well E6 with 20 ng/mL bFGF. Following 24-hours of FGF treatment, at which point cells were 100% confluent, cells were washed 1x with PBS and dissociated using 1.5 mL/well of Accutase in the incubator for ∼10 minutes. Dissociated cells were collected, washed, and spun at 1200 rpm for 2 minutes. For each well that was dissociated, cells were resuspended in 12 mL of E6 media containing 1x CET, 10 uM SB 435142 (SB), 20 ng/mL bFGF, and 3-5 uM CHIR99021 (CHIR). Note, the CHIR dose should be adjusted for each cell line; WTC-11 and derivative cell lines require 3 uM CHIR, while WTB and H9 require 5 uM CHIR. Cells were plated into Geltrex-coated 24-well plates using 0.8 mL of cell suspension per well. 48-hours later, cells had a cobble-stone appearance covering 100% of the well, and were either collected to assess NMP identity or immediately transitioned into day 5 differentiation media (see V2a differentiation below).

### V2a differentiation

The NEP V2a differentiation followed the detailed protocol outlined in Butts et al. (2019) with minor modifications. To start differentiation (day 0), stem cell cultures at ∼70% confluency were dissociated using Accutse, counted, and resuspended in mTeSR Plus with 1x CET, 10 uM SB, and 200 nM LDN 193189 (LDN) to a concentration of 100k cells/mL. 1 mL of cell suspension was plated into each well of Geltrex-coated 24-well plates, resulting in a final seeding density of ∼50k cells/cm^2^. At days 1 and 3, cells were fed with 2 mL/well mTeSR Plus with 10 uM SB and 200 nM LDN. By day 5, cells were 100% confluent. From day 5 onward, the base media was changed to Neural Induction Media (NIM; DMEM:F12 containing 1x N2 supplement, 1x Gluta-gro, 1x NEAA, 1x PenStrep, 0.4 ug/mL ascorbic acid, and 2 ug/mL heparin). On day 5, cells were fed with 2 mL/well NIM with 100 nM retinoic acid, 10 ng/mL BDNF, 10 uM SB, and 200 nM LDN. Day 5 also marks the time point at which NMPs are transitioned into the neural differentiation. From day 7 to 19, cells were fed every other day with 2 mL/well of NIM with 100 nM retinoic acid, 10 ng/mL BDNF, 0.4 ug/mL ascorbic acid, 300 nM purmorphamine, and 1 uM DAPT.

### V2a dissociation and cryopreservation

On day 19 of neuron differentiation, cultures were washed 1x with PBS then incubated for 20 min with 1 mL/well of Accutase. An additional 1 mL of PBS was then added to each well and neural cultures were broken up by smoothly pipetting up and down 10x. Dissociated cells were then pooled in a vessel containing an additional 24 mL of PBS (or an additional mL of PBS for every well being dissociated). Dissociated cells were distributed to 50 mL conicals and centrifuged at 1000 rpm for 3 minutes and supernatant was aspirated, being careful not to disturb the cell pellet. Cells were resuspended in 5-8 mL of NIM with 100 nM RA, 10 ng/mL BDNF, 300 nM PUR, 1 uM DAPT, and 1x CET. The cell suspension was passed through a 37 um filter and counted using an automated cell counter and Trypan dye. Each well of a 24-well plate should yield 3-4e6 cells. The cell concentration was then adjusted to ∼8-10e6 cells/mL by adding additional supplemented NIM. Cells were then mixed in a 1:1 ratio with cell preservation media (80% FBS, 20% DMSO) and distributed into labeled cryovials. Prior to transfer to LN2 for long-term storage, cryovials were chilled in a -80C freezer in CoolCell (Corning) or Mr. Frosty (Thermo Fisher) containers to keep the cooling rate to -1°C per minute.

### Flow Cytometry and FACS

Cells were dissociated as described above. Cells were spun for 2 min at 1200 RPM, then supernatant was aspirated and cells resuspended in 100 uL of PBS. Single cell suspensions were fixed using the FOXP3/Transcription factor kit (eBiosciences) following the manufacturer’s protocol. Cells in the Perm Buffer were stored at 4°C until use. For staining, the desired volume of cells was transferred to a clean 1.5 mL tube and washed 1x with 500 uL of PBS containing 0.1% BSA. Cells were spun in a fixed angle centrifuge at 400 g for 1.5 minutes, then tubes were rotated 180° and spun again. Supernatant was aspirated leaving ∼25 uL of liquid in the bottom of the tube to not disturb the cell pellet. Primary and secondary antibodies were diluted in FOXP3 kit Perm Buffer and cells were stained in 100 uL volume in the dark at room temp for 1 hr or overnight at 4°C. Cells were washed 2x with PBS/BSA between the primary and secondary antibody steps. For antibody product numbers and dilutions, see **Supplemental Table 1**. All flow cytometry was carried out on an Attune NxT. For live cell sorting, dissociated cells were resuspended in DMEM/F12 with 0.5 uM EDTA and 1x CET to a concentration of ∼10e6 cells/mL. Cells were immediately sorted on a FACS Aria Fusion into the same media.

### Immunostaining

For staining cells in plates, cells were fixed with 4% PFA at room temperature for 10 minutes, then washed 3x with PBS. Cells were blocked for 1hr at room temperature in PBS containing 5% normal donkey serum and 0.3% Triton X-100. Primary and secondary antibodies were diluted in PBS containing 1% BSA and 0.3% Triton X-100 and incubated at room temperature for 1 hr hour or overnight at 4°C, with 3x washes of PBS following each antibody incubation. For antibody product numbers and dilutions, see **Supplemental Table 1**.

### Nuclei Isolation for 10X Multiome

For multiome, all samples had been previously cryopreserved. Samples were thawed, washed with 10 mL of NIM, resuspended in 0.5 mL of NIM with CET and counted using Trypan Blue dye to assess viability. For sample pooling, 333k cells from three technical replicates were pooled, and 500k cells were used for subsequent nuclei isolation. Nuclei isolation followed 10X Genomics Demonstrated Protocol CG000365 Rev C: Nuclei Isolation for Single Cell Multiome ATAC + Gene Expression Sequencing including the DNase treatment step and using 0.8x lysis buffer strength. Nuclei were submitted to the UCSF Genomics CoLabs for library preparation and sequenced by the UCSF Institute for Human Genetics.

### Multiome Data Analysis

For each sample, multiome sequencing data was aligned to the Human GRCh38 reference genome (10X Genomics) containing custom reference sequences for integrated transgenes using CellRanger ARC (v2.0.2). Background RNA transcripts were then removed using CellBender (v0.1) before creation of a dual RNA/ATAC object and data analysis using Seurat (v4.4.0) and (Signac v1.9.0). Cells were filtered to have between 500 and 10,000 detected genes, between 1,000 and 50,000 RNA UMI, TSS enrichment score greater than 1, a nucleosome signal less than 2, and at least 1000 ATAC UMI. DoubletFinder (v2.0.3) was implemented only on RNA data to detect and remove presumptive doublets. RNA assay data was log normalized and scaled, while ATAC assay data was processed by latent semantic indexing using the RunTFIDF() and RunSVD() functions. Both assays were incorporated for UMAP visualization using the FindMultimodalNeighbors() function, and clustering was performed on RNA data with FindClusters() using the Louvain algorithm with multilevel refinement (algorithm = 2) and a resolution of 0.8. Clusters were annotated as described in the text by examining relative expression levels of marker genes. To facilitate balanced analysis, samples from matched cell lines (i.e., WTC-NEP and WTC-NMP) were subset to approximately the same number of cells. Subsetting included taking a random sample of a fixed number of cells as well as all cells in p2 and V2a clusters since they represented the major cell types of interest. Only after each sample was individually annotated and subset were samples merged using Harmony integration (theta = 1, sigma = 0.6). 12,020 cells were included in the subset and integrated object (NEP, NMP, and IND samples).

In the merged data, differentially expressed genes were identified using the FindMarkers() function using the Wilcoxon test (test.use = ‘wilcox’). Differentially expressed genes were tested for enriched gene ontology terms using the enrichGO() function in the clusterProfiler (v4.4.4) package. Differentially accessible ATAC peaks were identified using FindMarkers() function (test.use = ‘wilcox’ and min.pct = 0.1). The genes near the differentially expressed peaks were tested for enriched gene ontology terms using the GREAT tool (http://great.stanford.edu/public/html/). Motifs were assigned to ATAC peaks with the AddMotifs() function and using the JASPAR2020 motif database. Motif enrichment in ATAC regions was run using the FindMotifs() function and compared to a set of 50,000 GC-content matched background peaks identified using MatchRegionStats(). For label transfer, human fetal samples were downloaded from GEO (GSE171890) with the provided cell annotations. Each sample was subset to ventral spinal cord cell types then integrated into a single object in Seurat. NEP and NMP *in vitro* datasets were mapped onto the *in vivo* data using FindTransferAnchors() and TransferData() functions.

### Calcium Imaging

Freshly thawed d19 neurons were counted and replated at a density of 150,000 cells per well into geltrex coated 96-well plates in NIM supplemented with 100 nM retinoic acid, 10 ng/mL BDNF, 0.4 ug/mL ascorbic acid, 300 nM Purmorphamine, 1 uM DAPT, and 1x CET cocktail. After two days, an additional 1x volume supplemented BrainPhys media was added to each well. BrainPhys media (StemCell Technologies) was supplemented with 1:50 SM1 supplement, 1:100 CultureOne supplement, 1x PenStrep, 10 ng/mL BDNF, and 10 ng/mL GDNF. At indicated time points, media was fully removed and replaced with BrainPhys Imaging Optimized Medium with 1 ug/mL Fluo4-AM (Ion Biosciences) and 1:100 Pluronic F-127 (Ion Biosciences) and incubated for 1 hour. Media was then again completely replaced with warmed BrainPhys Imaging Optimized Medium alone and cells were imaged on an ImageXpress Confocal HT (Molecular Devices) equipped with stage heating and CO_2_ control (37°C, 5% CO_2_) at 10x magnification for 1 minute at a frame rate of 2 Hz. After calcium imaging, cells were fed with supplemented BrainPhys media and returned to the incubator for continued maturation.

Calcium imaging data was analyzed using custom written python code. Dense cultures and overlapping cells prevented easy masking on cell bodies for ROI, so a more agnostic approach was taken. All images in a single imaging session were collapsed into a maximum projection and then a global threshold was applied to create a mask of the recorded cell area. The cell mask was then divided into a 64×64 grid, with each square of the grid being treated as a single ROI (4,096 potential ROI). All pixels within the ROI which were included in the threshold mask were included in fluorescent intensity calculations. ROI with no pixels included in the threshold mask were not quantified.

### Induced Line Creation

The NGN2:LHX3 induced cell line was produced by modifying an existing plasmid for induced motor neurons: PB-TO-hNIL was a gift from iPSC Neurodegenerative Disease Initiative (iNDI) & Michael Ward (Addgene plasmid # 172113). PB-TO-hNIL was amplified in a PCR reaction to remove the ISL1 coding sequence using the forward primer ATGGAAGCCAGAGGCGAAC and reverse primer AGGTCCAGGGTTCTCCTCCACGTCTC using KOD Xtreme^TM^ Hotstart Polymerase (Millipore) then digested with DpnI (NEB) and recircularized using a T4 ligase (NEB) blunt end ligation with PNK (NEB). The resulting plasmid was transformed into DH5ɑ competent *E. coli* (NEB) for colony selection and miniprep purification using a QIAprep Spin Miniprep Kit (QIAGEN). Deletion of ISL1 was confirmed by separate restriction digests with DraIII, SmaI, and BbsI. For transfection, WTC11 stem cells were passaged at a density of 25k cells/well into a 24-well plate. The following day, the cells were fed with fresh media for 1 hour prior to transfection using 250 ng each of the NGN2:LHX3 plasmid and pEIF1a::Transposase (gifted by Dr. M. Ward), and 1 uL of Lipofectamine Stem (Invitrogen) prepared according to manufacturer’s instructions. The following day, the entire population of cells was passaged into T25 flasks and allowed to grow for another 24 hours prior to selection with 0.5 ug/mL puromycin. Purified cells were expanded and cryopreserved until future use.

### Induced Neuron Differentiation

Induced neuron differentiation protocols were based on previously described protocols for NGN2 and NGN2:ISL1:LHX3 neurons <u>(Fernandopulle et al., 2018)</u>. For induced neuron differentiation, stem cells were grown to near confluency on Geltrex coated plates before changing to induction media (NIM supplemented with 1 uM DAPT, 2 ug/mL doxycycline (dox), and listed concentrations of RA (most efficient for NGN2:LHX3 line is 1 uM RA). Cells were then fed every other day. On day 2, cells received NIM with 2 ug/mL dox, RA, and 10 ng/mL BDNF. From day 4 onward, the media formulation was the same as day 2, but without dox.

### RNA Isolation and qPCR

RNA was isolated using the QIAGEN RNeasy Mini Plus Kit according to manufacturer’s instructions. Whenever possible, cells were lysed directly in the culture vessel. Note, for highly dense V2a differentiations from day 9-19, up to 1 mL of RLT with BME was used per well of a 24-well plate. Cell lysate was transferred to 1.5 mL tubes and directly placed in a -80°C freezer until RNA extraction. QIAShredder tubes were used to homogenize samples prior to gDNA clean-up and RNA isolation. RNA was eluted in 50 uL of nuclease-free water (Ambion) and quantified via nanodrop. If samples had insufficient concentration (<60 ng/uL) or salt contaminants (260/230 ratios < 1.5), they were further processed using the Zymo RNA Clean and Concentrator Kit. If not used immediately, RNA was stored at -80°C. RNA was converted to cDNA for qPCR using the qScript cDNA SuperMix (Quanta Bio) according to manufacturer’s instructions. qCPR reactions were set up in 10 uL volumes in 384-well plates using 20 ng of cDNA per reaction, 2X PowerUp SYBR Green Master Mix, and 200 nM of each primer. Custom primer sequences were ordered from IDT as single-stranded DNA oligonucleotides. qPCR reactions were run on a QuantStudio 6 Flex following the manufacturer recommended standard cycling conditions for the PowerUp SYBR Green Master Mix.

### Cell Oracle

Cells were filtered to include only those with scATAC-seq peaks between 2,000 and 20,000. Cicero was used to extract cis-regulatory connections and identify co-accessible peaks (Pliner et al., 2018). We scanned these peaks for transcription factor motifs using the motif scan function from CellOracle (Kamimoto et al., 2023).

For the scRNA-seq data, we followed similar preprocessing steps to analyses from previous section, and then filtered down cells to p2 and V2a annotated cells. The remaining clusters were then integrated based on their NMP/NEP identity using the Combat function in Scanpy (Wolf et al., 2018). Genes were filtered to include the top 3,000 most variable genes. The base gene regulatory network (GRN) was generated through the CellOracle pipeline by combining ATAC-seq peaks with motif information, after which centrality scores for each node in the GRN were calculated. Pseudotime analysis was performed based on NEP and NMP lineages.

Desired perturbations were simulated using the generated GRNs and the CellOracle pipeline. Subsequently, the gradient of shifts and effects on developmental pseudotime were calculated. Pre- and post-perturbation gene expression matrices were exported, and gene set enrichment analysis was performed using the GSEApy package with the c2.cp.reactome category (database version 2023.2.Hs) from the Msigdb class (Fang et al., 2023). Enrichment was analyzed with the gp.gsea function, utilizing the signal-to-noise ratio metric for gene ranking and permutation testing on pre- and post-perturbation data.

### Lentiviral shRNA production and knockdown

Cloning and lentivirus production followed recommended protocols by the Broad Institute Genetic Perturbation Platform (https://portals.broadinstitute.org/gpp/public/). Briefly, the pLKO.1 cloning vector (a gift from David Root; Addgene plasmid # 10878) was digested with AgeI and EcoRI and gel purified. Oligos containing shRNA sequences against target genes (see **Supplemental Table 2;** IDT) were annealed to form double stranded inserts and ligated into the pLKO.1 backbone. Two shRNA different sequences per target gene were made, for a total of 10 unique plasmids. Plasmid DNA was isolated using ZymoPURE Plasmid Miniprep Kit (Zymo) and confirmed using long-read whole plasmid sequencing (Plasmidsaurus).

For lentivirus production, pLKO.1-shRNA plasmid, pCMV-dR8.91, and pMD2.G (a gift from Didier Trono; Addgene plasmid # 12259) plasmids were combined in a 9:8:1 ratio and transfected into HEK293T cells using Nanofect Transfection reagent (ALSTEM) following the manufacturer’s protocol. Media was changed the following day and collected at 48 and 72 hours post transfection. Viral supernatant was stored at -20°C until use. For WTC11 stem cell transduction, viral supernatant was added to stem cell media in a 1:20 ratio with 10 ug/mL polybrene (Millipore). After two days, stem cells were selected using 0.5 ug/mL puromycin for an additional two days before subsequent expansion. Upon differentiation to V2a neurons by NEP and NMP routes, cultures were additionally selected with puromycin between day 7 and 9 to ensure that the shRNA construct had not been silenced.

### Bulk RNA Sequencing and Analysis

Bulk RNA sequencing libraries were produced using the Mercurious™ High sensitivity BRB-seq kit (Alithea Genomics). For the High sensitivity BRB-seq kit, purified RNA was first isolated using the RNeasy Mini Plus kit (QIAGEN). Libraries were sequenced on a NovaSeq X (Illumina) by the UCSF CAT Core. Code for sequence quality control, alignment, and generation of count matrices followed Mercurious^TM^ High sensitivity BRB-seq kit recommendations and used fastQC (v0.11.5) and STAR (v2.7.11b) packages. Samples with greater than 2e6 total counts were kept for further analysis. Counts and metadata tables were combined into a single dataset using DESeq2 (v1.36.0). For all-sample comparison using PCA, VST-normalized counts were extracted and batch corrected using the limma (v3.54.2) removeBatchEffect() function. For heatmap plotting, the size-factor normalized counts were extracted and used. For differential expression testing, the counts matrices were subset to only include either NEP or NMP data before generating the DESeq object using a ‘ ∼ batch + KD_Gene’ design. The lists of differentially expressed genes were filtered to only include genes that crossed the significance threshold in 1 or 2 samples. For single sample GSEA scoring, raw counts matrices were TPM normalized prior to running removeBatchEffect(). This corrected TPM matrix was then ranked and analyzed using singscore (v1.19.1) with the differentially expressed genes between NEP and NMP V2a neurons from Multiome being used as input gene sets.

## References

Alshawaf, A. J., Viventi, S., Qiu, W., D’Abaco, G., Nayagam, B., Erlichster, M., Chana, G., Everall, I., Ivanusic, J., Skafidas, E., & Dottori, M. (2018). Phenotypic and Functional Characterization of Peripheral Sensory Neurons derived from Human Embryonic Stem Cells. Scientific Reports, 8(1), 603. 10.1038/s41598-017-19093-0

Ampatzis, K., Song, J., Ausborn, J., & El Manira, A. (2014). Separate Microcircuit Modules of Distinct V2a Interneurons and Motoneurons Control the Speed of Locomotion. Neuron, 83(4), 934–943. 10.1016/j.neuron.2014.07.018

Azim, E., Jiang, J., Alstermark, B., & Jessell, T. M. (2014). Skilled reaching relies on a V2a propriospinal internal copy circuit. Nature, 508(7496), Article 7496. 10.1038/nature13021

Bonner, J. F., & Steward, O. (2015). Repair of spinal cord injury with neuronal relays: From fetal grafts to neural stem cells. Brain Research, 1619, 115–123. 10.1016/j.brainres.2015.01.006

Bubnys, A., Kandel, H., Kao, L. M., Pfaff, D., & Tabansky, I. (2019). Hindbrain v2a neurons pattern rhythmic activity of motor neurons in a reticulospinal coculture. Frontiers in Neuroscience, 13, 1077. 10.3389/fnins.2019.01077

Butts, J. C., Iyer, N., White, N., Thompson, R., Sakiyama-Elbert, S., & McDevitt, T. C. (2019). V2a interneuron differentiation from mouse and human pluripotent stem cells. Nature Protocols, 14(11), Article 11. 10.1038/s41596-019-0203-1

Butts, J. C., McCreedy, D. A., Martinez-Vargas, J. A., Mendoza-Camacho, F. N., Hookway, T. A., Gifford, C. A., Taneja, P., Noble-Haeusslein, L., & McDevitt, T. C. (2017). Differentiation of V2a interneurons from human pluripotent stem cells. Proceedings of the National Academy of Sciences of the United States of America, 114(19), 4969–4974. 10.1073/pnas.1608254114

Cambray, N., & Wilson, V. (2007). Two distinct sources for a population of maturing axial progenitors. Development, 134(15), 2829–2840. 10.1242/dev.02877

Clovis, Y. M., Seo, S. Y., Kwon, J.-S., Rhee, J. C., Yeo, S., Lee, J. W., Lee, S., & Lee, S.-K. (2016). Chx10 Consolidates V2a Interneuron Identity through Two Distinct Gene Repression Modes. Cell Reports, 16(6), 1642–1652. 10.1016/j.celrep.2016.06.100

Crone, S. A., Viemari, J.-C., Droho, S., Mrejeru, A., Ramirez, J.-M., & Sharma, K. (2012). Irregular Breathing in Mice following Genetic Ablation of V2a Neurons. The Journal of Neuroscience, 32(23), 7895–7906. 10.1523/JNEUROSCI.0445-12.2012

Debrulle, S., Baudouin, C., Hidalgo-Figueroa, M., Pelosi, B., Francius, C., Rucchin, V., Ronellenfitch, K., Chow, R. L., Tissir, F., Lee, S.-K., & Clotman, F. (2019). Vsx1 and Chx10 paralogs sequentially secure V2 interneuron identity during spinal cord development. Cellular and Molecular Life Sciences. 10.1007/s00018-019-03408-7

Elder, N., Fattahi, F., McDevitt, T. C., & Zholudeva, L. V. (2022). Diseased, differentiated and difficult: Strategies for improved engineering of in vitro neurological systems. Frontiers in Cellular Neuroscience, 16. https://www.frontiersin.org/articles/10.3389/fncel.2022.962103

Fang, Z., Liu, X., and Peltz, G. (2023). GSEApy: a comprehensive package for performing gene set enrichment analysis in Python. Bioinformatics 39, btac757. 10.1093/bioinformatics/btac757.

Fattahi, F., Steinbeck, J. A., Kriks, S., Tchieu, J., Zimmer, B., Kishinevsky, S., Zeltner, N., Mica, Y., El-Nachef, W., Zhao, H., de Stanchina, E., Gershon, M. D., Grikscheit, T. C., Chen, S., & Studer, L. (2016). Deriving human ENS lineages for cell therapy and drug discovery in Hirschsprung disease. Nature, 531(7592), 105–109. 10.1038/nature16951

Fernandopulle, M. S., Prestil, R., Grunseich, C., Wang, C., Gan, L., & Ward, M. E. (2018). Transcription Factor–Mediated Differentiation of Human iPSCs into Neurons. Current Protocols in Cell Biology, 79(1), e51. 10.1002/cpcb.51

Gouti, M., Tsakiridis, A., Wymeersch, F. J., Huang, Y., Kleinjung, J., Wilson, V., & Briscoe, J. (2014). In vitro generation of neuromesodermal progenitors reveals distinct roles for wnt signalling in the specification of spinal cord and paraxial mesoderm identity. PLoS Biology, 12(8), e1001937. 10.1371/journal.pbio.1001937

Hayashi, M., Hinckley, C. A., Driscoll, S. P., Moore, N. J., Levine, A. J., Hilde, K. L., Sharma, K., & Pfaff, S. L. (2018). Graded arrays of spinal and supraspinal v2a interneuron subtypes underlie forelimb and hindlimb motor control. Neuron, 97(4), 869–884.e5. 10.1016/j.neuron.2018.01.023

Henrique, D., Abranches, E., Verrier, L., & Storey, K. G. (2015). Neuromesodermal progenitors and the making of the spinal cord. Development, 142(17), 2864–2875. 10.1242/dev.119768

Kamimoto, K., Stringa, B., Hoffmann, C. M., Jindal, K., Solnica-Krezel, L., & Morris, S. A. (2023). Dissecting cell identity via network inference and in silico gene perturbation. Nature, 614(7949), 742–751. 10.1038/s41586-022-05688-9

Kamimoto, K., Stringa, B., Hoffmann, C.M., Jindal, K., Solnica-Krezel, L., and Morris, S.A. (2023). Dissecting cell identity via network inference and in silico gene perturbation. Nature 614, 742–751. 10.1038/s41586-022-05688-9.

Kathe, C., Skinnider, M. A., Hutson, T. H., Regazzi, N., Gautier, M., Demesmaeker, R., Komi, S., Ceto, S., James, N. D., Cho, N., Baud, L., Galan, K., Matson, K. J. E., Rowald, A., Kim, K., Wang, R., Minassian, K., Prior, J. O., Asboth, L., … Courtine, G. (2022). The neurons that restore walking after paralysis. Nature, 611(7936), 540–547. 10.1038/s41586-022-05385-7

Lippmann, E. S., Williams, C. E., Ruhl, D. A., Estevez-Silva, M. C., Chapman, E. R., Coon, J. J., & Ashton, R. S. (2015). Deterministic HOX patterning in human pluripotent stem cell-derived neuroectoderm. Stem Cell Reports, 4(4), 632–644. 10.1016/j.stemcr.2015.02.018

Madisen, L., Zwingman, T. A., Sunkin, S. M., Oh, S. W., Zariwala, H. A., Gu, H., Ng, L. L., Palmiter, R. D., Hawrylycz, M. J., Jones, A. R., Lein, E. S., & Zeng, H. (2010). A robust and high-throughput Cre reporting and characterization system for the whole mouse brain. Nature neuroscience, 13(1), 133–140. 10.1038/nn.2467

Menelaou, E., & McLean, D. L. (2019). Hierarchical control of locomotion by distinct types of spinal V2a interneurons in zebrafish. Nature Communications, 10. 10.1038/s41467-019-12240-3

Menelaou, E., VanDunk, C., & McLean, D. L. (2014). Differences in Spinal V2a Neuron Morphology Reflect Their Recruitment Order During Swimming in Larval Zebrafish. The Journal of Comparative Neurology, 522(6), 1232–1248. 10.1002/cne.23465

Metzis, V., Steinhauser, S., Pakanavicius, E., Gouti, M., Stamataki, D., Ivanovitch, K., Watson, T., Rayon, T., Mousavy Gharavy, S. N., Lovell-Badge, R., Luscombe, N. M., & Briscoe, J. (2018). Nervous System Regionalization Entails Axial Allocation before Neural Differentiation. Cell, 175(4), 1105–1118.e17. 10.1016/j.cell.2018.09.040

Ozair, M. Z., Kintner, C., & Brivanlou, A. H. (2012). Neural induction and early patterning in vertebrates. Wiley Interdisciplinary Reviews. Developmental Biology, 2(4), 479. 10.1002/wdev.90

Philippidou, P., & Dasen, J. S. (2013). Hox genes: Choreographers in neural development, architects of circuit organization. Neuron, 80(1), 12–34. 10.1016/j.neuron.2013.09.020

Pliner, H.A., Packer, J.S., McFaline-Figueroa, J.L., Cusanovich, D.A., Daza, R.M., Aghamirzaie, D., Srivatsan, S., Qiu, X., Jackson, D., Minkina, A., et al. (2018). Cicero Predicts *cis*-Regulatory DNA Interactions from Single-Cell Chromatin Accessibility Data. Mol. Cell 71, 858–871.e8. 10.1016/j.molcel.2018.06.044.

Rayon, T., Maizels, R. J., Barrington, C., & Briscoe, J. (2021). Single-cell transcriptome profiling of the human developing spinal cord reveals a conserved genetic programme with human-specific features. Development, 148(15), dev199711. 10.1242/dev.199711

Samuel, R. M., Navickas, A., Maynard, A., Gaylord, E. A., Garcia, K., Bhat, S., Majd, H., Richter, M. N., Elder, N., Le, D., Nguyen, P., Shibata, B., Llabata, M. L., Selleri, L., Laird, D. J., Darmanis, S., Goodarzi, H., & Fattahi, F. (2023). Generation of Schwann cell derived melanocytes from hPSCs identifies pro-metastatic factors in melanoma. BioRxiv. 10.1101/2023.03.06.531220

Shi, Y., Lin, S., Staats, K. A., Li, Y., Chang, W.-H., Hung, S.-T., Hendricks, E., Linares, G. R., Wang, Y., Son, E. Y., et al. (2018). Haploinsufficiency leads to neurodegeneration in C9ORF72 ALS/FTD human induced motor neurons. Nat. Med. 24, 313–325. doi:10.1038/nm.4490.

Squair, J. W., Milano, M., de Coucy, A., Gautier, M., Skinnider, M. A., James, N. D., Cho, N., Lasne, A., Kathe, C., Hutson, T. H., Ceto, S., Baud, L., Galan, K., Aureli, V., Laskaratos, A., Barraud, Q., Deming, T. J., Kohman, R. E., Schneider, B. L., … Anderson, M. A. (2023). Recovery of walking after paralysis by regenerating characterized neurons to their natural target region. Science, 381(6664), 1338–1345. 10.1126/science.adi6412

Tian, R., Gachechiladze, M. A., Ludwig, C. H., Laurie, M. T., Hong, J. Y., Nathaniel, D., Prabhu, A. V., Fernandopulle, M. S., Patel, R., Abshari, M., Ward, M. E., & Kampmann, M. (2019). CRISPR Interference-Based Platform for Multimodal Genetic Screens in Human iPSC-Derived Neurons. Neuron, 104(2), 239–255.e12. 10.1016/j.neuron.2019.07.014

Tzouanacou, E., Wegener, A., Wymeersch, F. J., Wilson, V., & Nicolas, J.-F. (2009). Redefining the Progression of Lineage Segregations during Mammalian Embryogenesis by Clonal Analysis. Developmental Cell, 17(3), 365–376. 10.1016/j.devcel.2009.08.002

Wang, C., Ward, M. E., Chen, R., Liu, K., Tracy, T. E., Chen, X., Xie, M., Sohn, P. D., Ludwig, C., Meyer-Franke, A., Karch, C. M., Ding, S., & Gan, L. (2017). Scalable Production of iPSC-Derived Human Neurons to Identify Tau-Lowering Compounds by High-Content Screening. Stem Cell Reports, 9(4), 1221–1233. 10.1016/j.stemcr.2017.08.019

Wolf, F.A., Angerer, P., and Theis, F.J. (2018). SCANPY: large-scale single-cell gene expression data analysis. Genome Biol. 19, 15. 10.1186/s13059-017-1382-0.

Wymeersch, F. J., Skylaki, S., Huang, Y., Watson, J. A., Economou, C., Marek-Johnston, C., Tomlinson, S. R., & Wilson, V. (2019). Transcriptionally dynamic progenitor populations organised around a stable niche drive axial patterning. Development, 146(1). 10.1242/dev.168161

Zhang, Y., Pak, C., Han, Y., Ahlenius, H., Zhang, Z., Chanda, S., Marro, S., Patzke, C., Acuna, C., Covy, J., Xu, W., Yang, N., Danko, T., Chen, L., Wernig, M., & Südhof, T. C. (2013). Rapid Single-Step Induction of Functional Neurons from Human Pluripotent Stem Cells. Neuron, 78(5), 785–798. 10.1016/j.neuron.2013.05.029

Zholudeva, L. V., & Lane, M. A. (2019). Choosing the right cell for spinal cord repair. Journal of Neuroscience Research, 97(2), 109–111. 10.1002/jnr.24351

Zholudeva, L. V., Iyer, N., Qiang, L., Spruance, V. M., Randelman, M. L., White, N. W., Bezdudnaya, T., Fischer, I., Sakiyama-Elbert, S. E., & Lane, M. A. (2018). Transplantation of Neural Progenitors and V2a Interneurons after Spinal Cord Injury. Journal of Neurotrauma, 35(24), 2883–2903. 10.1089/neu.2017.5439

Zholudeva, L. V., Karliner, J. S., Dougherty, K. J., & Lane, M. A. (2017). Anatomical Recruitment of Spinal V2a Interneurons into Phrenic Motor Circuitry after High Cervical Spinal Cord Injury. Journal of Neurotrauma, 34(21), 3058–3065. 10.1089/neu.2017.5045

